# Proteases as Biological Bits for Programmable Medicine

**DOI:** 10.1101/607895

**Authors:** Brandon Alexander Holt, Gabriel A. Kwong

**Affiliations:** Wallace H. Coulter Department of Biomedical Engineering, Georgia Tech College of Engineering and Emory School of Medicine, Atlanta, GA 30332, USA; Parker H. Petit Institute of Bioengineering and Bioscience, Atlanta, GA 30332, USA; Institute for Electronics and Nanotechnology, Georgia Tech, Atlanta, GA 30332; Integrated Cancer Research Center, Georgia Tech, Atlanta, GA 30332; The Georgia Immunoengineering Consortium, Emory University and Georgia Tech, Atlanta, GA 30332

## Abstract

Engineered biocircuits that interface with living systems as plug-and-play constructs may enable new applications for programmable therapies and diagnostics. We create biological bits (bbits) using proteases – a family of pleiotropic, promiscuous enzymes – to construct the biological equivalent of Boolean logic gates, comparators and analog-to-digital converters. We use these modules to write a cell-free bioprogram that can combine with bacteria-infected blood, quantify infection burden, and then calculate and unlock a selective drug dose. Inspired by probabilistic computing, we leverage multi- and common-target protease promiscuity as the biological analog of superposition to program three probabilistic bbits that solve all implementations of the two-bit oracle problem, Learning Parity with Noise. Treating a network of dysregulated proteases in a living animal as an oracle, we use this algorithm to resolve the probability distribution of coagulation proteases *in vivo*, allowing diagnosis of pulmonary embolism with high sensitivity and specificity (AUROC = 0.92) in a mouse model of thrombosis. Our results demonstrate that protease activity can be programmed in cell-free systems to carry out classical and probabilistic algorithms for programmable medicine.

## Introduction

Rapid advances in engineered biological circuits are motivating the design of new treatment and detection platforms for practical applications in programmable medicine. The development of foundational components, such as molecular logic gates^1^ and genetic clocks^2,3^, have enabled the design of biocircuits with increasing complexity, including the ability to solve mathematical problems^4^, build autonomous robots^5^, and play interactive games^6^. Recently, programmable biocircuits have been applied for therapeutic and diagnostic applications^7^, including genetic circuits that sense-and-respond to dysregulated inflammation^8^ or blood glucose levels^9^. To date, the design of these biocircuits is principally focused on constructs that are implemented in cell-based platforms – which require genome or protein engineering^10–13^ – and carry out algorithms inspired by classical computer circuits, which operate on binary digits (bits) and Boolean logic gates (e.g., AND, OR, NAND).

While classical biocircuits are well-suited at performing deterministic tasks (e.g., input determines output)^14^, the ability to perform inference-based tasks – such as identification of which single input cause resulted in the observed output effect given multiple plausible inputs – are more challenging. In contrast to classical circuits, probabilistic circuits, which operate on analog bits characterized by a probability distribution of states, efficiently solve inference problems by assigning a likelihood probability that each plausible input would produce the observed output^15^. Probabilistic bits have been implemented with magnets (p-bits)^16–18^ as well as photons and electrons in quantum systems (qubits)^19,20^. In medicine, differential diagnoses are fundamentally based on inference, wherein an observed symptom could be caused by several diseases. Conversely, the decision to treat a patient is determined by a clear set of inputs (e.g., disease stage, biomarker level, etc.)^7^. For these reasons, we sought to develop a unified system of biological bits capable of executing both classical and probabilistic algorithms for therapeutic and diagnostic applications.

In living organisms, high-level functions arise from intricate networks of enzymatic activity that ultimately control complex systems ranging from immunity to blood homeostasis^21–24^ Among enzymes, proteases are both ubiquitous, comprising 2% of the human genome^25^, and promiscuous, having the ability to cleave diverse substrate sequences (6–8 AA) in addition to their putative target^26–33^. To leverage these features for programmable medicine, we define protease activity acting on a target substrate as a biological bit (bbit). Under a classical framework, a register of bbits comprise distinct protease-substrate pairs that take on the binary state 1 above an activity threshold (**Fig. 1A left., Fig. S1A**). By contrast, probabilistic bbits are constructed using promiscuous proteases that act on two substrates simultaneously to create a state of superposition where the probability of being measured in state 0 or 1 is based on relative substrate cleavage velocities (v0 and v1) (**Fig. 1A right, Fig. S1B**). Here we use classical bbits to design a plug-and-play therapeutic biocircuit capable of quantifying input bacterial activity and outputting a digital drug dose to clear infected human blood (**Fig 1B**). Under a probabilistic framework, we construct diagnostic biocircuits using probability-based gates to first solve the oracle problem Learning Parity with Noise (LPN), and then extend this system *in vivo* by treating networks of dysregulated proteases as a hidden oracle to noninvasively diagnose pulmonary embolism with high accuracy in a mouse model of thrombosis (**Fig 1C**).

**Figure 1.**
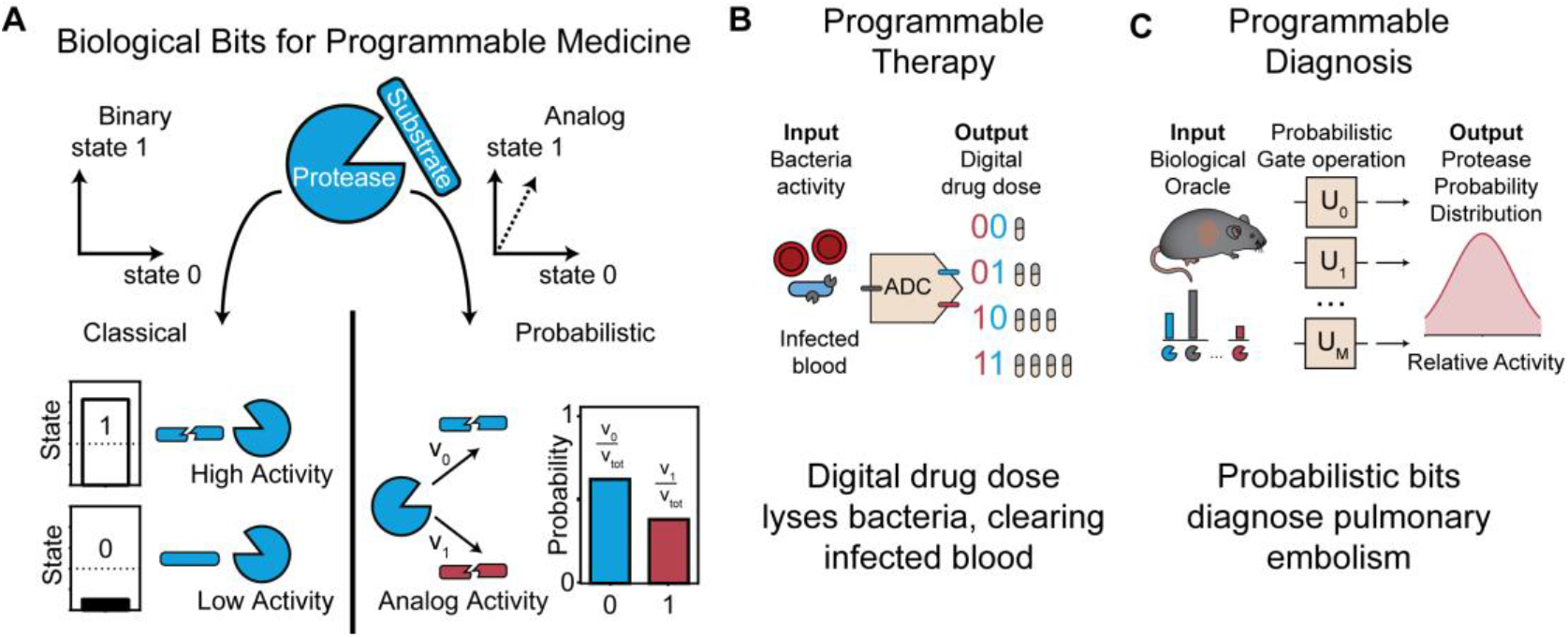
Protease activity as classical or probabilistic biological bits for programmable medicine. **(A)** (**left**) The binary state of a classical bit represented as two orthogonal states (0 or 1). A classical bbit exists in a state of either high or low protease activity, defined by a threshold (dotted line). (**right**) The binary state of probabilistic bits represented as a superposed vector between state 0 and state 1. A probabilistic bbit acting on two state substrates has two cleavage velocities (v_0_ and v_1_), which are the probabilities of observing the bbit in either state (0 or 1). **(B)** Binary biological bits are applied to construct a therapeutic biocircuit for digital drug delivery to clear infected blood of bacteria. **(C)** Probabilistic biological bits are applied to construct a noninvasive diagnostic that detects pulmonary embolism in living mice.

## Results

A central function of complex circuits is the ability to store and manipulate digitized information; therefore, we first set out to construct a flash analog-to-digital converter (ADC) to convert continuous biological signals into binary digits. An electronic ADC performs three major operations during signal conversion: voltage comparison, priority assignment, and digital encoding. An analog input voltage is first compared against a set of increasing reference voltages (V_0_–V_i_) by individual comparators (d_0_–d_i_) that allows current to pass if the input signal is greater than or equal to its reference value (**Fig. S2**). During priority assignment, only the activated comparator with the highest reference voltage, dn, remains on while all other activated comparators, d_n-1_–d_0_ are turned off. The prioritized signal is then fed into a digital encoder comprising OR gates to produce binary values. To design an ADC biocircuit using protease activity as the core signal, we constructed biological analogs of comparators by using liposomes locked by an outer peptide cage^34,35^ (**Fig. 2A; Fig. S3A, B**). With increasing peptide crosslinking densities, these biocomparators (b_0_–b_i_) served to reference the level of input protease activity (GzmB) required to fully degrade the peptide cage (IEFDSGK, **Table S1**) and expose the lipid core (**Fig. S3C**), analogous to the reference voltages stored in electronic comparators. We used lipase^36^ as a Buffer gate to open all biocomparators with fully degraded cages (**Fig. 2B, C; Fig. S4**) and release a unique combination of inhibitors and signal proteases (WNV, TEV, and WNV inhibitor) that collectively act to assign priority to the highest activated biocomparator (b_n_) by inhibiting all signal proteases released from other biocomparators (b_0_–b_n-1_). To encode the prioritized signal into binary values, we designed a set of OR gates using orthogonal quenched substrates (RTKR and ENLYFQG) specific for the signal proteases (WNV and TEV respectively; **Fig. 2D**) to provide fluorescent 2-bit readouts (p_0_–p_i_; **Fig. S5**). Fully integrated, our 4-2 bit biological ADC converted input protease levels (GzmB) across four orders of magnitude into binary digital outputs (**Fig. 2E, F**).

**Figure 2.**
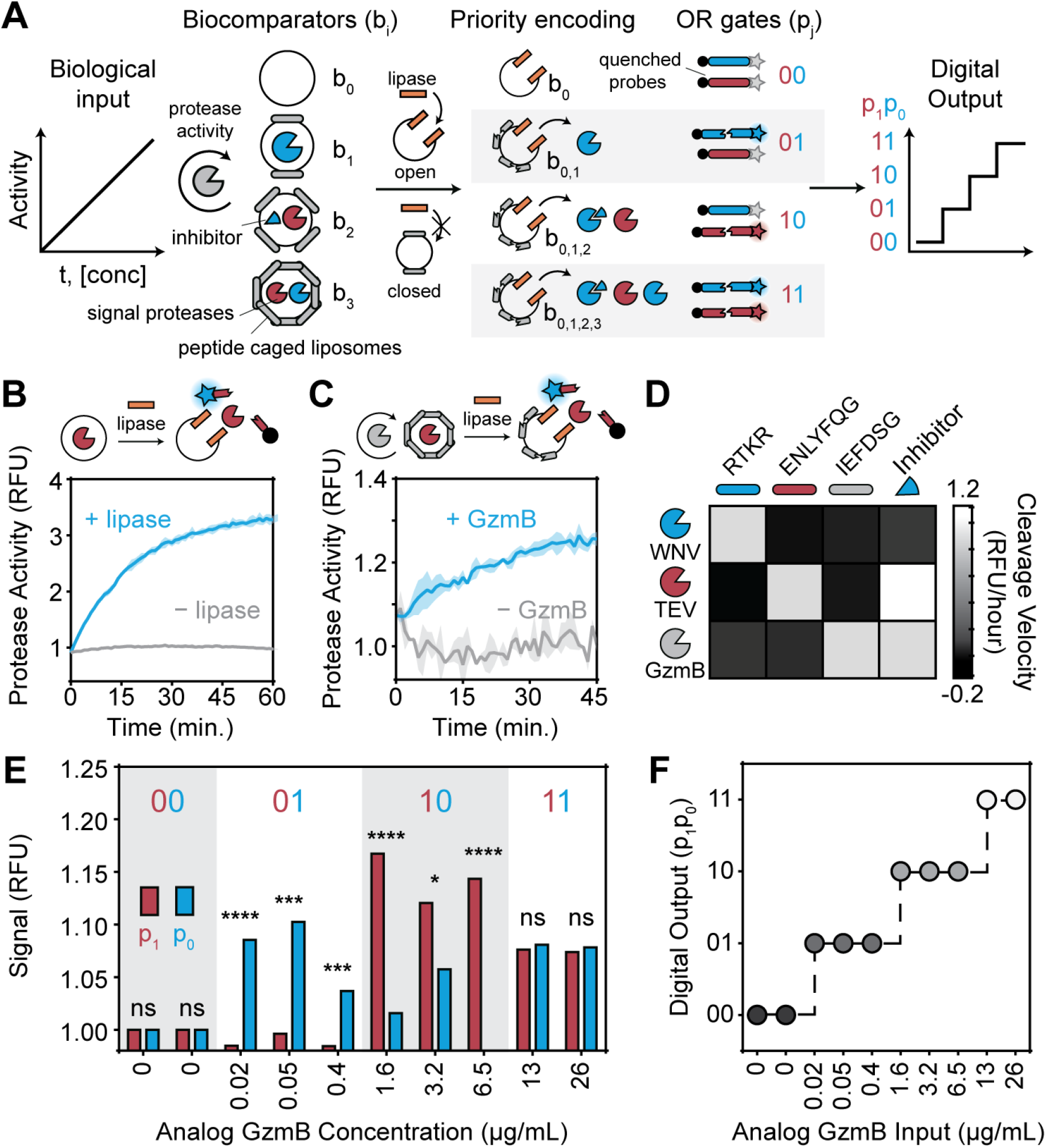
A biological ADC transduces protease activity to classical bits. (**A**) Biocircuit diagram depicting the conversion of a biological input (protease activity) into a digital output with biocomparators, buffer gates, and OR gates. Blue triangle inhibits blue protease activity. Circular arrow represents enzyme activity. (**B**) Bare or (**C**) peptide-caged liposomes opened by lipase or GzmB activity, respectively, release TEV protease that cleaves a quenched peptide substrate. Standard deviation represented by line shading. (**D**) Protease cleavage velocity orthogonality map measuring GzmB, WNV & TEV protease activity against respective substrates alone and in the presence of WNV protease inhibitor. (**E**) Concentrations of GzmB across four orders of magnitude are input to the bioADC, and bbits p_0_ and p_1_ are read out in a fluorescent assay (n = 3, paired two-way t-test). (**F**) Digital output as a function of input GzmB concentration. * < 0.05, ** < 0.01, *** < 0.001, and **** < 0.0001.

To demonstrate a practical biomedical application, we next sought to interface our biological ADC with a living system as a plug-and-play therapeutic biocircuit for digital drug delivery. We rewired our ADC to autonomously quantify input bacterial activity and then output an anti-microbial drug dose to selectively clear infected red blood cells (RBCs) of bacteria (DH5α *Escherichia coli)* (**Fig. 3A**). To construct biocomparators with the ability to prioritize input levels of bacterial activity, we synthesized liposomes with peptide cages using a substrate (RRSRRVK) specific for the *E. Coli* surface protease OmpT^37,38^ (**Fig. 3B**). We synthesized a series of 8 biocomparators with increasing peptide densities (0–10.2 nM) and validated their ability to sense input bacterial concentrations across 8 log units (0–10^8^ CFU/ml) using a fluorescent reporter (**Fig. S6**). To convert the release of signal proteases to a drug output, we designed protease-activatable prodrugs comprising cationic (polyarginine) anti-microbial peptides (AMP) (**Fig. 3C, Table S1**) in charge complexation with anionic peptide locks (polyglutamic acid)^39^ to block the activity of AMP. These drug-lock peptides were linked in tandem by OR gate peptides p_0_ and p_1_ (RTKR and ENLYFQG respectively) to allow signal proteases that directly cleave p_0_ or p_1_ to digitally control the output drug dose (**Fig. 2**). We designed one-third and two-thirds of the total drug dose to be unlocked by cleavage of p_0_ and p_1_, respectively, such that binary values 00, 01, 10, and 11 corresponded to 0/3, 1/3, 2/3, and 3/3 of the total drug dose (**Fig. S7**).

**Figure 3.**
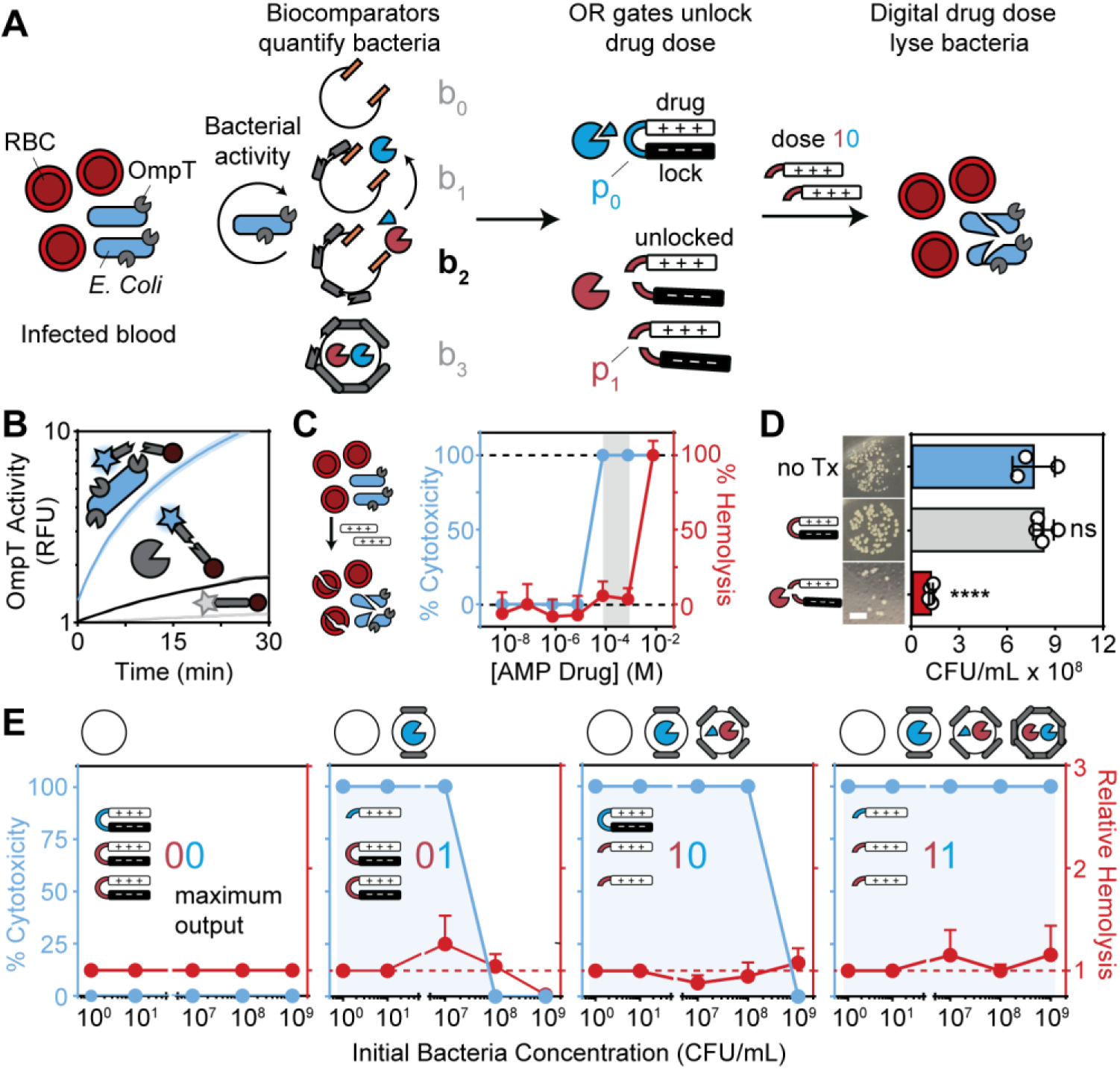
A fully integrated bioADC to execute an antimicrobial program. (**A**) Biocircuit depicting the use of an ADC to quantify bacteria and autonomously unlock digital drug doses. Circular arrow represents enzyme activity. (**B**) Cleavage assay measuring recombinant OmpT and live *E. coli* culture cleavage of peptide RRSRRV (n = 3, two-way ANOVA & Sidak’s multiple comparisons). (**C**) EC50 measurement for drug cytotoxicity and hemolysis against *E. coli* and RBCs, respectively. Gray shading represents therapeutic window with 100% cytotoxicity and no hemolysis. (**D**) Viability of bacteria after treatment with locked drug and locked drug + protease. (n = 4, one-way ANOVA & Dunnett’s multiple comparisons to bacteria only control; scale bar = 4 mm). (**E**) Drug bacteria cytotoxicity and RBC hemolysis at five concentrations of bacteria with 4 different versions of the antimicrobial program each containing a different number of biocomparators (two-way ANOVA & Sidak’s multiple comparisons; hemolysis n = 2, cytotoxicity n = 3). Line shading and error bars are standard deviation. * < 0.05, ** < 0.01, *** < 0.001, and **** < 0.0001.

To confirm the therapeutic efficacy of our prodrug design, treatment of bacteria with locked drug had no significant cytolytic activity compared to untreated controls, but by contrast, treatment with protease-cleaved drug-lock complexes resulted in a significant reduction in bacterial colonies (**Fig. 3D**). We observed similar levels of bacterial cytotoxicity when AMP was directly loaded into liposomes, showing that charge complexation was required to fully block AMP activity (**Fig. S6B**). In human RBCs infected with *E. coli* at concentrations ranging from 10^0^–10^9^ CFU/mL, samples containing a single biocomparator (b0) lacked the ability to eliminate bacteria as anticipated (output = 00). By contrast, increasing the number of biocomparators in the samples (b_0_–b_3_) allowed our program to autonomously increase the drug dose (output 01, 10, and 11) in response to higher bacterial loads to completely eliminate infection burdens across 9 orders of magnitude up to 10^9^ CFU/mL without significantly increasing hemolysis (**Fig. 3E, Fig. S8**). Our data showed that cell-free biocircuits can be constructed using protease activity as a primary digital signal to execute autonomous drug delivery programs under a broad range of conditions.

Our antimicrobial ADC used protease activity as binary classical bbits to carry out a sense-and-respond bioprogram for drug delivery. To demonstrate the use of protease activity as probabilistic bbits to solve inference problems, we designed probabilistic circuits by leveraging protease promiscuity. Protease promiscuity occurs under multi-target (i.e., a single protease cutting multiple substrates) and common-target (i.e., multiple proteases cutting the same substrate) settings, and is a fundamental feature that allows proteases to carry out distinct physiological functions^26^ (e.g., coagulation proteases control the formation of fibrin clots as well as the expression of adhesion molecules and cytokines^40^) (**Fig S9E**). To create two-state probabilistic bbits, we considered the superposed activity of a single protease cleaving two distinct substrates, and defined the probability of the protease to be found in state 0 (cleaving substrate 1) or state 1 (cleaving substrate 2) by the relative cleavage velocity for either substrate^41^ (**Fig. 4A; Fig. S9**). This allowed state probabilities (i.e., cleavage velocities) to be quantitatively controlled according to Michaelis-Menten^41^ kinetics by changing the substrate concentration or sequence.

**Figure 4.**
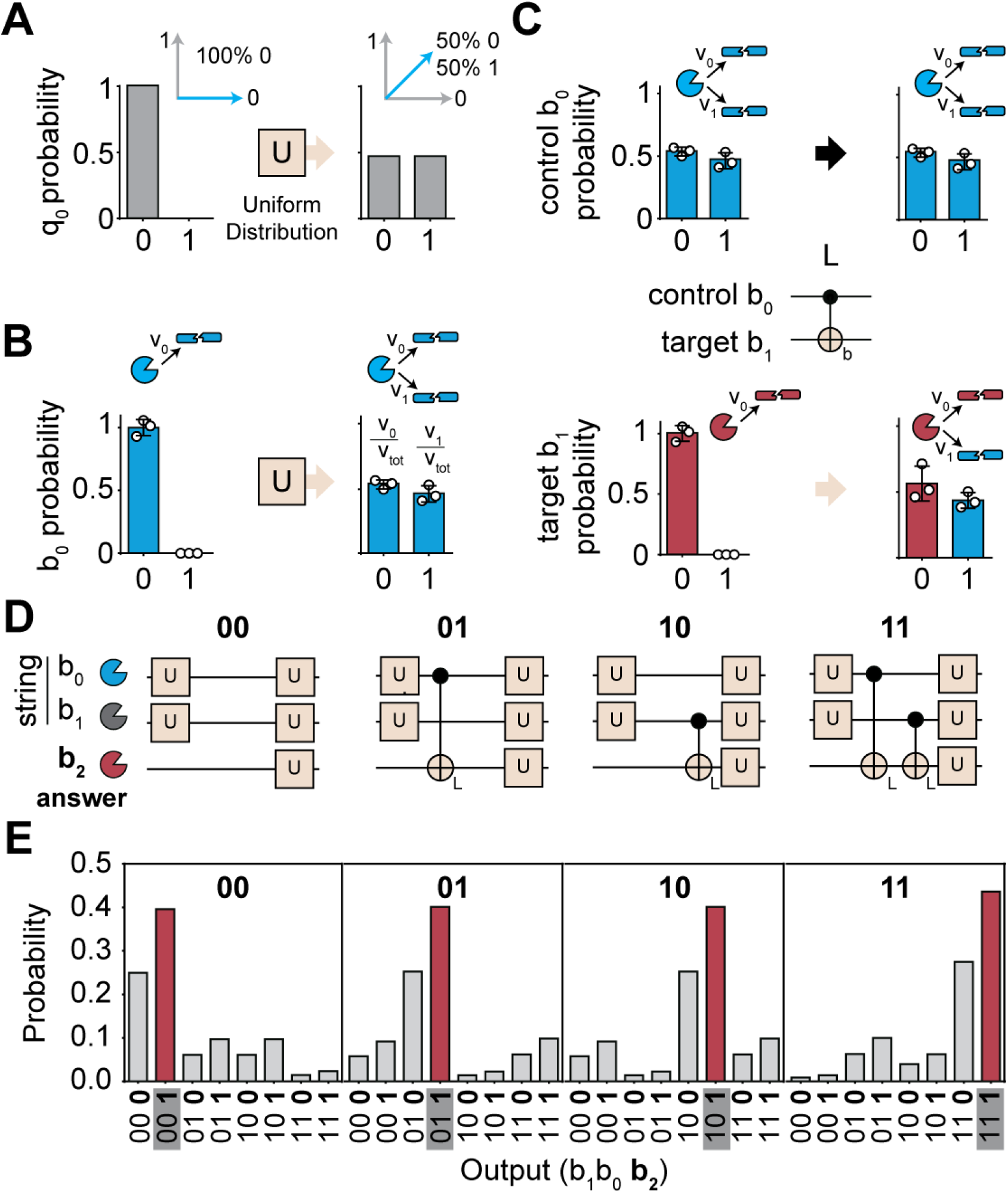
Superposing bbits to solve oracle-based problems with probabilistic logic gates. (**A**) Ideal inputs and outputs of a Uniform (U) gate acting on the basis state 0. (**B**) Implementation of the biological U gate. A protease (thrombin), first only exposed to the state 0 substrate, is then exposed to the state 1 substrate, resulting in similar probabilities of being observed in either state. (**C**) Control protease bit (thrombin) and target protease bit (plasmin) share similar specificity for the same state 1 substrate such that the probability of each protease cutting in the 1 state is correlated. (**D**) Biological scores represent four configurations of the 2-bit oracle problem. (**E**) Bbits solve all four implementations of the two-bit oracle problem.

Under this framework for quantifying protease probabilities, we built a set of biological probabilistic gates to perform operations on state probabilities that we named the Uniform gate (U-gate) and Linker gate (L-gate). These gates make use of multi- and common-target promiscuity, and we designed their operations based on previous implementations of probabilistic gates to solve a classic oracle problem, Learning Parity with Noise (LPN)^42^, where the goal is to deduce the value of a 2-bit string hidden by the oracle in the fewest number of calls (**Fig S9, Table S1, 2**). Analogous to conducting a coin flip, we designed our U-gate to create a superposition of states by taking an input bbit, b_0_, in state 0 with 100% probability (i.e., single substrate cleavage) and outputting b_0_ in state 0 or 1 with equal probability (i.e., performed by adding a second substrate to allow multi-target cleavage) (**Fig. 4A, B**). By contrast, analogous to a classical XOR gate, we designed the L-gate to take two input bbits – control and target bbits b_0_ and b_1_ respectively – and operate on the state 1 probability of target b_1_ such that it exhibits parity, or is linked, to the state 1 probability of control b_0_ (i.e., performed by adding a second substrate to allow common-target cleavage between two proteases) (**Fig. 4C, Fig S9E**). We constructed biological scores, based on probabilistic scores^42^, to implement all four instances of the 2-bit LPN problem by using our U- and L-gates to operate on three protease bits – 2 string bits (b_0_ and b_1_) to represent possible hidden string values (00, 01, 10, and 11) and 1 answer bit (b_2_) (**Fig. S9B, C, Table S1, 2**). By multiplying all permutations of the output state 0 and 1 probabilities of bbits b_0_–b_2_ (**Fig. S9D**), our protease solver correctly deduced the value of the hidden string among all other possibilities by assigning it the highest probability in all four oracle configurations (**Fig. 4E; Fig. S9D**). Collectively, our results showed that protease activity can be quantified as state probabilities and operated by probabilistic logic gates to efficiently solve inference problems.

We next sought to demonstrate a practical application using probabilistic bbits as a diagnostic platform for disease detection in living animals. The ability to detect dysregulated protease activity has important diagnostic applications for broad diseases, such as the prothrombin time (PTT) assay which is used to diagnose thrombosis^43^. Here we considered dysregulated protease networks, such as those in thrombosis, to be represented as an oracle string of protease activities with a distinct probability distribution compared to a healthy state (**Fig. 5A**). Analogous to the Central Limit Theorem (CLT), we postulated that designing promiscuous, common-target substrates to detect dysregulated protease networks could be modeled as sampling a probability distribution where sampling means would converge to normal distributions even if the underlying protease probability distribution is itself not normally distributed. We therefore sought to design and adapt a new set of U-gates to sample differences in uniformity between dysregulated and healthy protease networks, and use the resulting normalized variances (σ^2^) to discriminate disease (**Fig 5A**).

**Figure 5.**
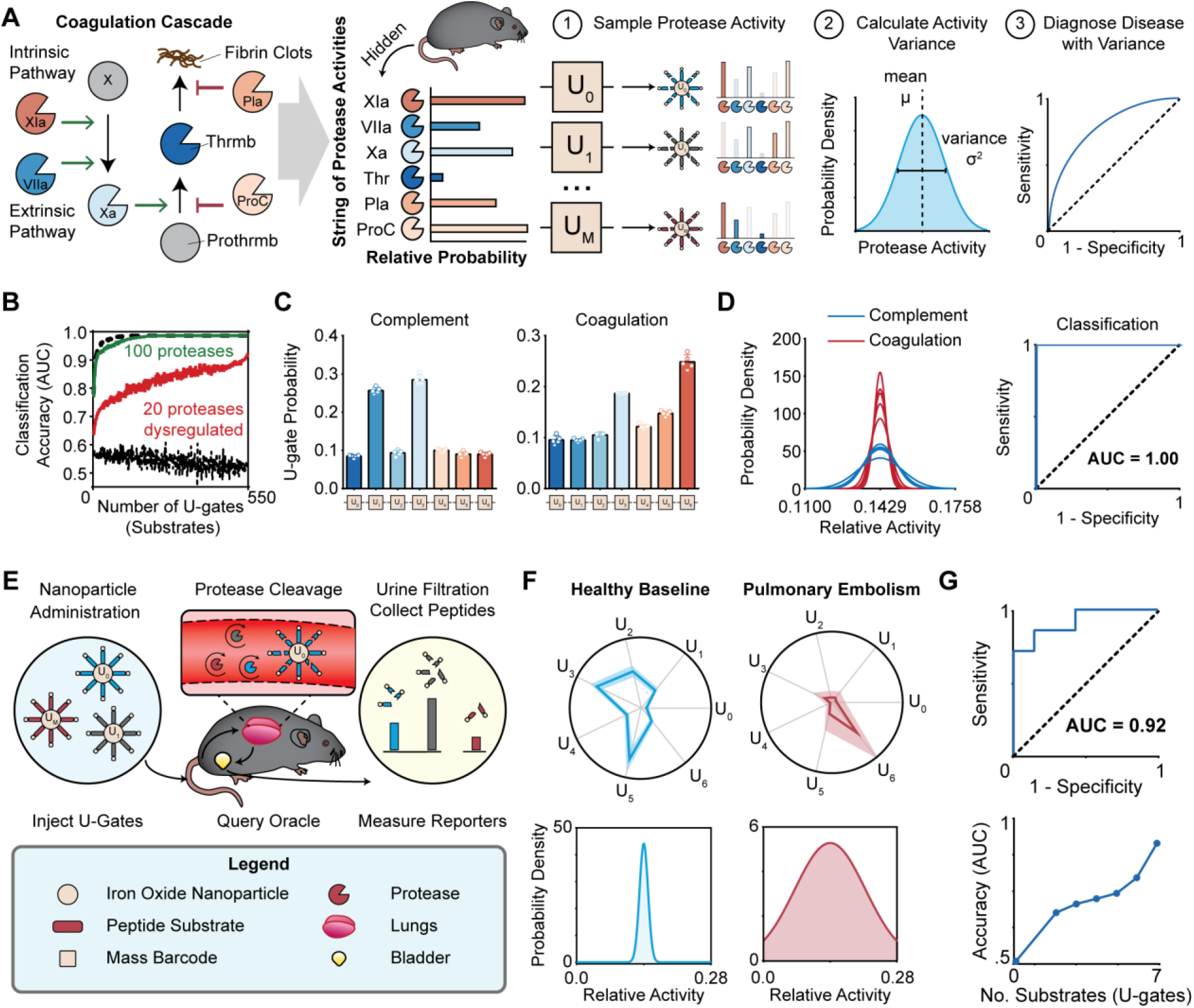
Applying the biological oracle algorithm to noninvasively detect pulmonary embolism. **(A)** Coagulation cascade comprised of protease network can form or degrade fibrin clots. The individual protease activities can be represented as a string of numbers, by supposing that the organism behaves as a biological oracle. The relative activity of each protease is sampled when multiple proteases cut the same substrate (i.e., U-gate). The variance of the U-gate signals reflects the uniformity of the underlying string of protease activities (b_0_—b_N_). The variance is used to classify mice as diseased or healthy. **(B)** Simulation of healthy and diseased protease networks, where variable numbers of proteases are dysregulated in the disease case. Measuring the effect of increasing number of substrates on the disease classification accuracy, given variable numbers of dysregulated proteases. Bottom dashed black line represents control case where no proteases are dysregulated, and top dashed black line represents control case were *all* 550 proteases are dysregulated. **(C)** Activity signatures of complement (left) and coagulation (right) protease cocktails against the seven U-gates (U_0_—U_6_). Error bars represent standard deviation (n = 5). **(D)** (left) Measuring variance from all coagulation and complement mixtures, plotted as the normal probability distribution functions. (right) Using the U-gate variance to classify the protease cocktails as complement or coagulation. **(E)** Nanoparticles carrying peptide substrates (U-gates) are injected in to the mouse and proteases cleave off mass tag reporters from each gate. The reporters are filtered into the urine and collected for quantification by mass spectrometry. **(F)** (top) Urinalysis measuring the concentration of each of the seven U-gates on Day 0 (healthy) and Day 4 (disease) averaged between all seven animals. Maximum quantity for healthy and disease plots is 0.1 and 0.3, respectively. Shading is standard deviation. (bottom) Measuring variance from one example healthy and disease mouse, plotted as the normal probability distribution function. **(G)** (top) Classifying pulmonary embolism with U-gate variance. (bottom) Classification accuracy increases with increased number of substrates (plotted as mean of all possible combinations).

To test this approach computationally, we randomly generated baseline activity scores between zero and one for all 550 proteases encoded in the human genome^26^. Random strings of 0, 20, 100, or 550 proteases were upregulated or downregulated in equal proportion by scaling their activity by a factor of five to reflect an average of literature reported values^44–46^. To simulate promiscuous sampling by a set of U-gates, we modeled a substrate library of size *M* (ranging from 2–550) randomly sampling *n* proteases (ranging from 1–550) by adding corresponding activity scores and computing the probability distribution and normalized variance across all U-gates. The results from our model revealed that the ability to classify disease and healthy networks increases as the number of dysregulated proteases (red and green traces; **Fig. 5B**) or U-gates increases (e.g., greater than 90% classification accuracy can be achieved with >10 U-gates and >20 dysregulated proteases) (**Fig 5B**). This result showing dependence of classification accuracy on feature size was consistent with computational results based on multidimensional datasets^47^. To validate our model prediction, we designed seven substrates (U_0_–U_6_) to sense the complement (e.g., C1r, MASP2, Factor D, Factor I) and coagulation protease networks (e.g., thrombin, plasmin, factor XIIa, factor Xa, protein C) (**Fig S10**). Using the measured U-gate outputs after incubation with either group of proteases *in vitro* (**Fig 5C**), the normalized variances of the U-gate outputs classified mixtures as either complement or coagulation with perfect accuracy (n = 10, AUROC = 1.00, **Fig 5D**). These results confirmed that a set of promiscuous U-gates can be used to sample and discriminate differences in the underlying probability distributions of protease networks.

To apply this approach for *in vivo* diagnostics, we used a thromboplastin-induced mouse model of pulmonary embolism (PE)^48^ to test whether our library of U-gates could discriminate mice with blood clots from healthy controls. Recently, we developed a class of protease activity sensors for delivery of mass-barcoded peptide substrates to quantify protease activity *in vivo*^28,49,50^. Mass-barcoded substrate libraries are conjugated to a nanoparticle carrier, delivered intravenously, and upon protease cleavage, release substrate fragments that are cleared into urine for quantification by mass spectrometry according to their mass barcode. Using this platform, we administered a single cocktail of our seven mass-barcoded U-gates to quantify protease activity in healthy mice (**Fig. 5E; Fig. S11**)^28^ as well as in mice induced with PE (**Fig. 5F**). The measured variance across our 7 U-gates noninvasively diagnosed PE with high sensitivity and specificity (AUROC = 0.92) (**Fig. 5F, G; Fig. S12**), and consistent with our mathematical predictions, overall classification accuracy increased from 0.5 to 0.92 as the number of U-gates used in the classifier increased from zero to seven respectively (**Fig. 5G**). Collectively, our data showed that by treating strings of protease activity as a probability distribution, the underlying sample variance can be used to infer and diagnose pulmonary embolism with high accuracy.

## Discussion

By interpreting protease activity as carrying binary or probabilistic information, we demonstrated the use of proteases as biological bits in cell-free biocircuits for therapeutic and diagnostic applications. We used the classical interpretation of protease bbits to construct a 2-bit analog-to-digital converter (ADC) as an autonomous drug delivery biocircuit to clear infected blood of bacteria across 9 orders magnitude in concentration. To construct our biological ADC, we designed biocomparators using peptide-caged liposomes because these materials are well-tolerated and biologically compatible^51,52^. Our cell-free approach is distinct from cell-based genetic circuits^7,53^ that require significant protein or organismal engineering to control signaling, including the non-trivial OFF state for proteases which has required insertion strategies^11–13^ artificial autoinhibitors^31^, or dimerizing leucine zippers^53^ to control. Cell-free liposomes have also been used in past studies such as synthetic minimal cells to control the expression of genetic circuits by liposome fusion^54,55^. Our approach may be amenable to integration with these genetic approaches, if for example, these circuits were redesigned to input or output proteases.

In contrast to classical binary bits, we also explored the use of protease activity as probabilistic bits. By leveraging both multi-target and common-target protease promiscuity, we designed logic gates to operate on the probability states of protease bbits to provide the ability to solve inference-based oracle problems, such as LPN, by deducing the correct value of hidden strings with the highest probability. We further considered dysregulated protease networks within a living animal as representing a biological oracle with a distinct probability distribution of states, which enabled us to noninvasively diagnose thrombosis with high classification accuracy. This approach is similar to the Central Limit Theorem (CLT) where sampling an unknown probability distribution, regardless whether the probability distribution itself is normally distributed or not, will result in sample means that converge to a normal distribution with variance proportional to the unknown distribution^56^. Therefore, we designed promiscuous U-gates to sample the underlying probability distribution of dysregulated protease networks such as the coagulation cascade, and using as few as seven substrates, achieved a disease classification accuracy > 0.92 *in vivo*. As there are >15 proteases involved in these cascades, we envision that the future use of larger (>50) substrate libraries may allow development of pan-diagnostics capable of monitoring whole-organism protease activities (>250 extracellular proteases).

Under both classical and probabilistic frameworks, our biological circuits were designed to sense extracellular proteases, which we do not envision will limit potential *in vitro* or *in vivo* applications. Of the greater than 550 proteases encoded by the genome, over half are secreted or membrane-bound and involved in a host of different diseases^26,57^. A significantly greater diversity of secreted and membrane bound proteases is represented by bacterial and viral species^58–62^. This rich diversity has provided the biological foundation for biomedical applications that rely on extracellular protease activation of pro-drug or pro-diagnostics in living animals including patients^57,58^. In our work, we provided examples of multiple types of biocircuits (ADC, logic gates, comparators, etc.) that are modular and can be engineered to input or output bacterial (OmpT), viral (TEV, WNV), murine (coagulation cascade), or mammalian (GzmB) proteases in both *in vitro* and *in vivo* settings. Looking forward, by integrating the full richness of protease biology and promiscuity, harnessing proteases as binary or probabilistic bits may provide a unique biological advantage for programmable control of future therapeutics and diagnostics.

## Acknowledgments

This work was funded by an NIH Director’s New Innovator Award (Award No. DP2HD091793). B.A.H is supported by the NSF GRFP, National Institutes of Health GT BioMAT Training Grant under Award Number 5T32EB006343 and the Georgia Tech President’s Fellowship. This material is based upon work supported by the National Science Foundation Graduate Research Fellowship under Grant No. **DGE-1650044** (B.A.H.). G.A.K. holds a Career Award at the Scientific Interface from the Burroughs Welcome Fund. We acknowledge Dr. D. Myers (Georgia Tech and Emory) and S. N. Dahotre (Georgia Tech and Emory) for their helpful discussions during the preparation of the manuscript. The authors thank Quoc Mac (Georgia Tech) for assistance with the mouse model. The content is solely the responsibility of the authors and does not necessarily represent the official views of the National Institutes of Health. We acknowledge use of the IBM Q for this work. The views expressed are those of the authors and do not reflect the official policy or position of IBM or the IBM Q team.

## Author contributions

B.A.H. and G.A.K. designed research; B.A.H. performed research; B.A.H. and G.A.K. analyzed data; and B.A.H and G.A.K. wrote the paper.

## Data and materials availability

All data is available in the main text or the supplementary materials.

## Supplementary Materials

### Materials and Methods

#### 1 Animals

6-to 8-week old female mice were used at the outset of all experiments. Mice were purchased from Charles River Laboratories. All animal protocols were approved by Georgia Tech IACUC (protocol #A17097).

#### 2 Protease Cleavage Assays

All protease cleavage assays were performed with a BioTek Cytation 5 Imaging Plate Reader, taking fluorescent measurements at 485/528 nm and 540/575 nm (excitation/emission) for read-outs measuring peptide substrates terminated with FITC (Fluorescein isothiocyanate) and 5-TAMRA (5-Carboxytetramethylrhodamine), respectively. Kinetic measurements were taken every minute over the course of 60 – 120 minutes at 37 C. West Nile Virus NS3 protease (WNVp) and Tobacco Etch Virus protease (TEVp), along with their substrates, inhibitors and buffers were obtained from Anaspec, Inc. (Fremont, CA). Phospholipase C (PLC), Phosphatidylinositol-Specific (from Bacillus cereus) was purchased from Thermo Fisher Scientific (Waltham, MA). Activity RFU measurements were normalized to time 0 measurement, and as such represent fold change in signal. Granzyme B (GzmB) was purchased from PeproTech, Inc. (Rocky Hill, NJ). Thrombin and Factor XIa were purchased from Haematologic Technologies (Essex, VT). Outer Membrane Protease T (OmpT, Protease 7) was purchased from Lifespan Biosciences (Seattle, WA). C1r was purchased from Millipore Sigma (Burlington, MA). GzmB, Thrombin, Factor XIa, and C1r fluorescent peptide substrates were custom ordered from CPC Scientific (Sunnyvale, CA). OmpT fluorescent peptide substrate was custom ordered from Genscript (Piscataway, NJ). See **Table S1** and **Table S2** for more information regarding proteases, substrates, and inhibitors.

2.1 Figure 2B 10 uL of liposomes (34 mM) loaded with TEVp (1 ug protease/17 mmol liposome) were coincubated with 50 uL TEVp substrate in provided activity buffer (pH 7.5). 2 uL of PLC (100 units/mL) was added to the experimental group, and 2 uL of assay buffer was added to the control group.
2.2 Figure 2C 10 uL of liposomes (34 mM) loaded with TEVp (1 ug protease/17 mmol liposome), embedded with 10 mol% CPAA and crosslinked at 0.1% efficiency with GzmB substrate were coincubated with 50 uL TEVp substrate in provided activity buffer (pH 7.5). 2 uL PLC (100 U/mL) was added to both the control and experimental group. 2 uL GzmB (0.1 ug/uL) was added only to experimental group.
2.3 Figure 2D All amounts of protease, substrate, and inhibitor for WNVp and TEVp were added according to instructions from Anaspec WNVp and TEVp activity kit. All conditions incubated with WNVp inhibitor include protease of interest incubated with its primary substrate. GzmB was added at a working concentration of (0.01 mg/mL) to 2 uM of its peptide substrate.
2.4 Figure 2E All 4 biocomparator levels (b0-b3, 50 mM each) were added together (10 uL each), and co-incubated with 13 uL of GzmB solution (concentration varies depending on condition), 2 uL of PLC (100 U/mL), 0.5 uL of WNVp substrate (after diluted 100X according to manufacturer’s instructions), 0.5 uL of TEVp substrate (after diluted 100X according to manufacturer’s instructions), and 4 uL of assay buffer. Biocomparator levels 0-3 are referenced by peptide cage crosslinking efficiencies of 0, 0.01, 1 and 100%, respectively. Plotted values are taken at minute 30 and normalized to starting values (time 0, or equivalently, the no protease control). Paired t-tests were also performed between the Q0 and Q1 digit for each condition.
2.5 Figure 3B For recombinant OmpT condition, 2 uL of OmpT (0.5 mg/mL) was added to 18 uL of 2 uM OmpT substrate. For E. coli condition, 2 uL of E. coli (10^9^ CFU/mL) was added to 18 uL of 2 uM OmpT substrate. 2 uL of DI H2o was added to negative control, along with 18 uL of OmpT substrate.
2.6 Figure 4B and 4C 2 uL of Thrombin (10.1 mg/mL) or Plasmin (6.9 mg/mL) were added to 18 uL of respective state 1 or state 0 substrate (2 uM). Velocities of Thrombin and Plasmin cutting their respective state 1 or state 0 substrates were calculated over the first 5 minutes of incubation. These velocities were normalized by the sum of the state 1 and state 0 velocity. Therefore, the values plotted as probability in Figure 4 are relative velocities.
2.7 Figure 4E Protease bbit configurations for each implementation of the Oracle are referenced in **Fig. S10D**, and in each case 2 uL of protease was added to 2 uM of respective state substrate. H gates are made reversible by adding in the original state substrate (state-0) at a concentration 10-fold the new state (state-1). Stock concentrations of the proteases involved were: Factor XIa (6 mg/mL), Plasmin (6.9 mg/mL), Thrombin (10.1 mg/mL), and C1r (1 mg/mL). State 0 and State 1 peptide substrates were CC1 and CC6 for FXIa, CC4 and CC1 for C1r, CC2 and CC6 for Thrombin, and CC2 and CC9 for Plasmin. Probability of two digit state is calculated by multiplying the probability (relative velocity) for each individual protease bbit. For example, if the first digit (bbit) is Protease A, with relative velocities V_a1_ and V_a2_, and the second digit is Protease B, with velocities V_b1_ and V_b0_, then the probability of achieving the answer 01 = V_b0_*V_a1_.

#### 3 Liposome Synthesis and Characterization

Liposome synthesis kit, PIPES buffer, EDC*MeI, and spin filters (100 kDa m.w.c.o.) were purchased from Millipore Sigma (Burlington, MA). Cholesterol-anchored Polyacrylic Acid (4400 g/mol, 30-40 COOH groups/molecule, structure in **Fig. S3A**) was custom ordered from Nanocs (Boston, MA). Float-a-lyzer dialysis tubes (100 kDa m.w.c.o., 1 mL) were purchased from Spectrum Labs (Rancho Dominguez, CA). Synthesis protocol is adapted from the methods used by Basel et. al. (29). Liposomes were loaded with respective protease inhibitor cocktail amounts, and concentration was estimated via absorbance. Standard curve for estimating concentration of liposomes was used by correlating absorbance of liposome solution at 230 nm with known standard concentrations (**Fig. S3B**). CPAA was vortexed in warm water (< 10 mg/mL) and volume was added such that there was 10 mol% CPAA relative to the molarity of lipids in the liposome solution. Solution was incubated for 1 hour at room temperature, or overnight at 4 C. Excess polymer and materials were removed via centrifugation (spin filters, 3-5 times at 4700 XG for 10 mins) or float-a-lyzer membranes (4C in spinning water overnight). EDC*MeI was dissolved into 10 mM PIPES buffer and volume was added such that EDC*MeI:CPAA ratio was 4:1. Solution was incubated for 20 minutes at room temperature. Excess EDC was filtered out via centrifugation or dialysis tubes. Peptide crosslinker was added at desired molar ratio and incubated for 1 hour at room temperature or 4 C. Excess peptide was filtered via centrifugation or dialysis tubes. Change in liposome hydrodynamic diameter was measured via DLS on a Zetasizer Nano ZS, Malvern Panalytical (Netherlands). Volumes loaded into biocomparators include concentrations of proteases and inhibitors as follows: b_0_ = empty; b_1_ = 20 uL WNVp (0.1 mg/mL) + 80 uL DI H_2_O; b_2_ = 50 uL WNV inhibitor (1 uM) + 50 uL TEVp (0.04 mg/mL); b3 = 50 uL WNVp (0.1 mg/mL) + 50 uL TEVp (0.08 mg/mL).

#### 4 Bacterial Cytotoxicity & Human Red Blood Cell Hemolysis Assays

##### Bacterial culture and cytotoxicity measurement

DH5α *Escherichia coli* were a gift from Todd Sulchek’s BioMEMS lab at Georgia Tech. E. coli were cultured in LB broth (Lennox) at 37 C and plated on LB agar (Lennox) plates. LB broth was purchasd from Millipore Sigma (Burlington, MA) and LB agar was purchased from Invitrogen (Carlsbad, CA). AMP and locked AMP were custom ordered from Genscript (Piscataway, NJ). See **Table S1** for more information. Bacteria were grown to a concentration of 10^9^ CFU/mL before being used for experiments. Concentration was estimated by measuring the OD_600_ of the bacterial suspension, and assuming an OD_600_ of 1.000 corresponds to a concentration of 8 × 10^8^ CFU/mL. Bacterial cell viability was measured by making eight 10-fold serial dilutions, and plating three 10 uL spots on an LB agar plate. Plates were incubated overnight at 37C, and CFUs were counted. Untreated bacteria CFU counts served as control for 0% cytotoxicity, and bacteria + IPA (or 0 countable CFUs) served as control for 100% cytotoxicity.

##### RBC collection and hemolysis measurement

Healthy blood donors had abstained from aspirin in the last two weeks, and consent was obtained according to GT IRB H15258. Blood was drawn by median cubital venipuncture into sodium citrate (3.2%). The sample was subsequently centrifuged at 150 G for 15 min, and the resulting platelet rich plasma was discarded. Red blood cells were then washed three times with phosphate buffered saline (PBS). For each wash, 12 mL of PBS were added, the sample was centrifuged at 1000 RPM for 10 min, and the supernatant was discarded. Hemolysis was estimated by spinning down experimental RBC samples and measuring the absorbance of the supernatant at 450 nm. Absorbance values corresponding to 100% hemolysis came from incubating RBCs with 0.1% Tween-20. Absorbances corresponding to 0% hemolysis came from untreated RBCs.

4.1 Figure 3C For bacterial cytotoxicity measurements, 25 uL of antimicrobial peptide (AMP) was added, pertaining to 7 concentrations ranging between 7.6 nM and 7.6 mM. 20 uL of bacteria (10^7^ CFU/mL) were added, and the sample was filled to 200 uL with LB broth in PCR tubes. Sample tubes were taped on a plate shaker (250 RPM) incubating at 37 C for 8 hours. For RBC hemolysis measurements, the same assay was performed, but used 20 uL of donor RBCs instead of bacteria solution.
4.2 Figure 3D For bacteria only condition, 5 uL of bacteria (10^9^ CFU/mL) were added to 95 uL LB broth. For bacteria + AMP p_1_, 58 uL of AMP p_1_ (1.7 mM) were added to 5 uL of bacteria (10^9^ CFU/mL), with the solution being filled to 100 uL with LB broth. For bacteria + protease + locked AMP p_1_, 20 uL of TEVp (4 ug/mL) and 58 uL of AMP p_1_ (1.7 mM) were added to 5 uL of bacteria (10^9^ CFU/mL), with the solution being filled to 100 uL with LB broth. Samples in PCR tubes were taped to a plate shaker (250 RPM) incubating at 37 C for 1 hour. Serial dilutions and plating were then performed to measure viable bacteria concentrations.
4.3 Figure 3E Each condition includes 20 uL of the bioprogram (2 uL of PLC, 6 uL D1, 6 uL D2, 6 uL D3), 20 uL of bacteria, 10 uL of RBCs, 24 uL of locked peptide drug (9 uL of 1.7 mM AMP p_1_ and 15 uL of 0.53 mM AMP p_0_), and 126 uL PBS. The concentration of bacteria, and the presence of each biocomparator, depends on the experimental condition (**Fig. 3E**). Samples in PCR tubes were taped to a plate shaker (250 RPM) incubating at 37 C for 8 hours, followed by dilutions/plating to estimate bacterial cytotoxicity. The remainder of the sample was spun down by centrifugation and used to estimate hemolysis.

#### 5 Nanosensor synthesis and characterization

Aminated IONPs were synthesized in house per published protocol^28^. Mass barcode-labelled substrate peptides synthesized by MIT Core Facility and used for in vivo formulation. Aminated IONPs were first reacted to the heterobifunctional crosslinker Succinimidyl Iodoacetate (SIA; Thermo) for 2 hours at room temperature (RT) and excess SIA were removed by buffer exchange using Amicon spin filter (30 kDa, Millipore). Sulfhydryl-terminated peptides and Polyethylene Glycol (PEG; LaysanBio, M-SH-20K) were mixed with NP-SIA (90:20:1 molar ratio) and reacted overnight at RT in the dark to obtain fully conjugated activity nanosensors. Activity nanosensors were purified on a Superdex 200 Increase 10-300 GL column using AKTA Pure FPLC System (GE Health Care). Ratios of FITC per IONP were determined using absorbance of FITC (488 nm, ε = 78,000 cm^−1^M^−1^) and IONP (400 nm, ε = 2.07 × 10^6^ cm^−1^M^−1^)^35,81^ measured with Cytation 5 Plate Reader (Biotek). At this conjugation condition, our resulting formulations have an average of 50 FITC-labelled peptides per nanoparticle core. DLS measurements of activity nanosensors were done in PBS or mouse plasma at RT using Zetasizer Nano ZS (Malvern).

#### 6 Urinary Prediction of Blood Clots in Murine Model of Pulmonary Embolism

All urinalysis experiments were done in paired setup. Before (4 days prior) onset of thrombosis, mice were administered with peptide substrate-labelled activity nanosensors (50 ug of IONP per animal). Mice were placed over 96-well polystyrene plates surrounded by an open cylindrical sleeve covered by a weighted petri dish to prevent animals from leaving the cylinder. Thrombosis, or pulmonary embolism, was initiated by coinjecting 1.75 ug/g b.w and 0.17 mg/mouse of fibrinogen (0.5 nmol). of rabbit thromboplastin. Animals were left to urinate for 30 minutes before urine samples were collected. Individual substrates were quantified by mass spectrometry, which was performed as a service by Syneos Health.

#### 7 Statistical Analysis

Statistical analysis was performed using statistical packages included in GraphPad Prism 6. To assess the significance of increase in signal due to protease cleavage, we used a two-way ANOVA (without repeated measures) followed by Sidak’s multiple comparisons test (**Fig. 2B, C**, and **3B**). To assess the accuracy of assigning the binary value 0 or 1 to the digits p_0_ and p_1_ as seen in **Fig. 2E**, two-way paired t-tests were performed between the signal value from each digit to determine if the signal from one was statistical more prominent than the other (**Fig. 2E, 4B, C**). A one-way ANOVA followed by Dunnett’s multiple comparisons test was used to compare experimental means to cells only control in **Fig. 3D.** Two-way ANOVA followed by Sidak’s multiple comparisons test used to compare experimental means to control for bacterial cytotoxicity and RBC hemolysis (**Fig. 3E**).

**Fig. S1.**
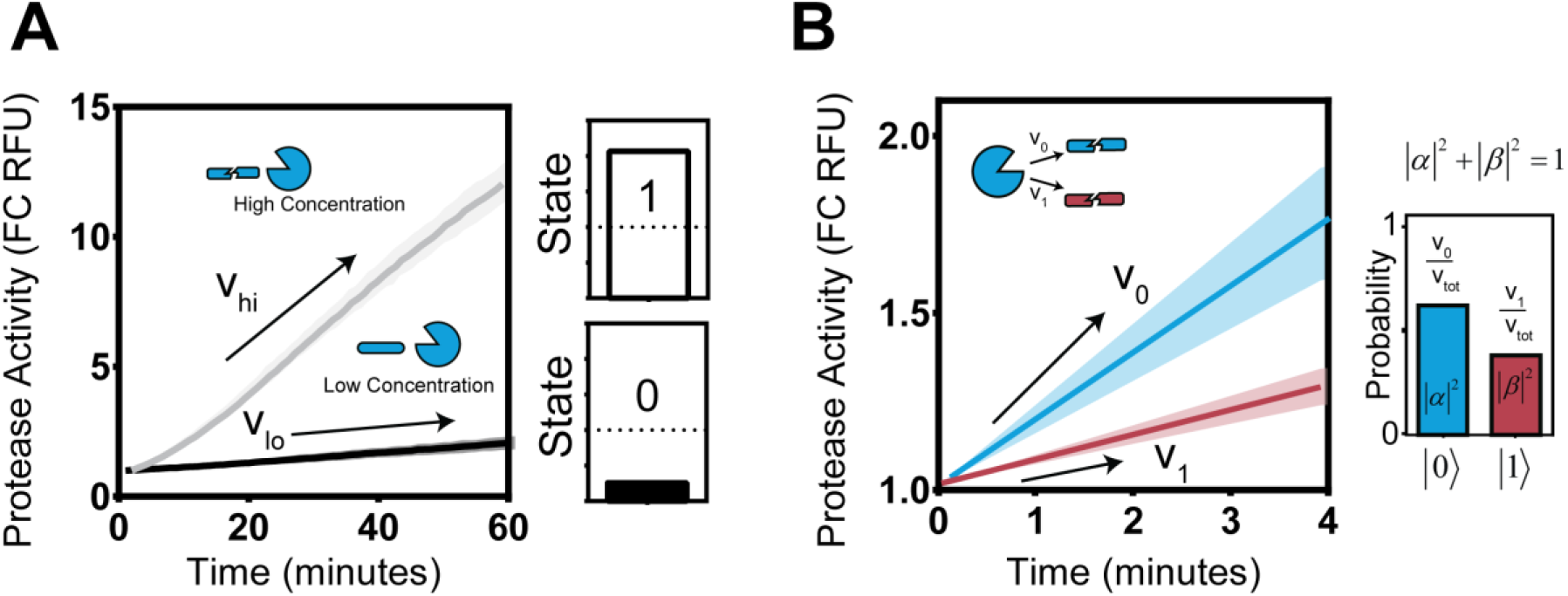
Extracting bit values from protease cleavage velocity. **(A)** Protease (C1r) activity assay against substrate (LQRIYK), at high and low activities (controlled with concentrations of protease). Velocity threshold is defined to classically separate state 1 (high activity) from state 0 (low activity). **(B)** Protease (Plasmin) activity assay against substrate state 0 (GLQRALEI) and state 1 (KYLGRSYKV). Relative velocities represent the probability of observing the protease cutting in either state. Line shading represents standard deviation (n = 3).

**Fig. S2.**
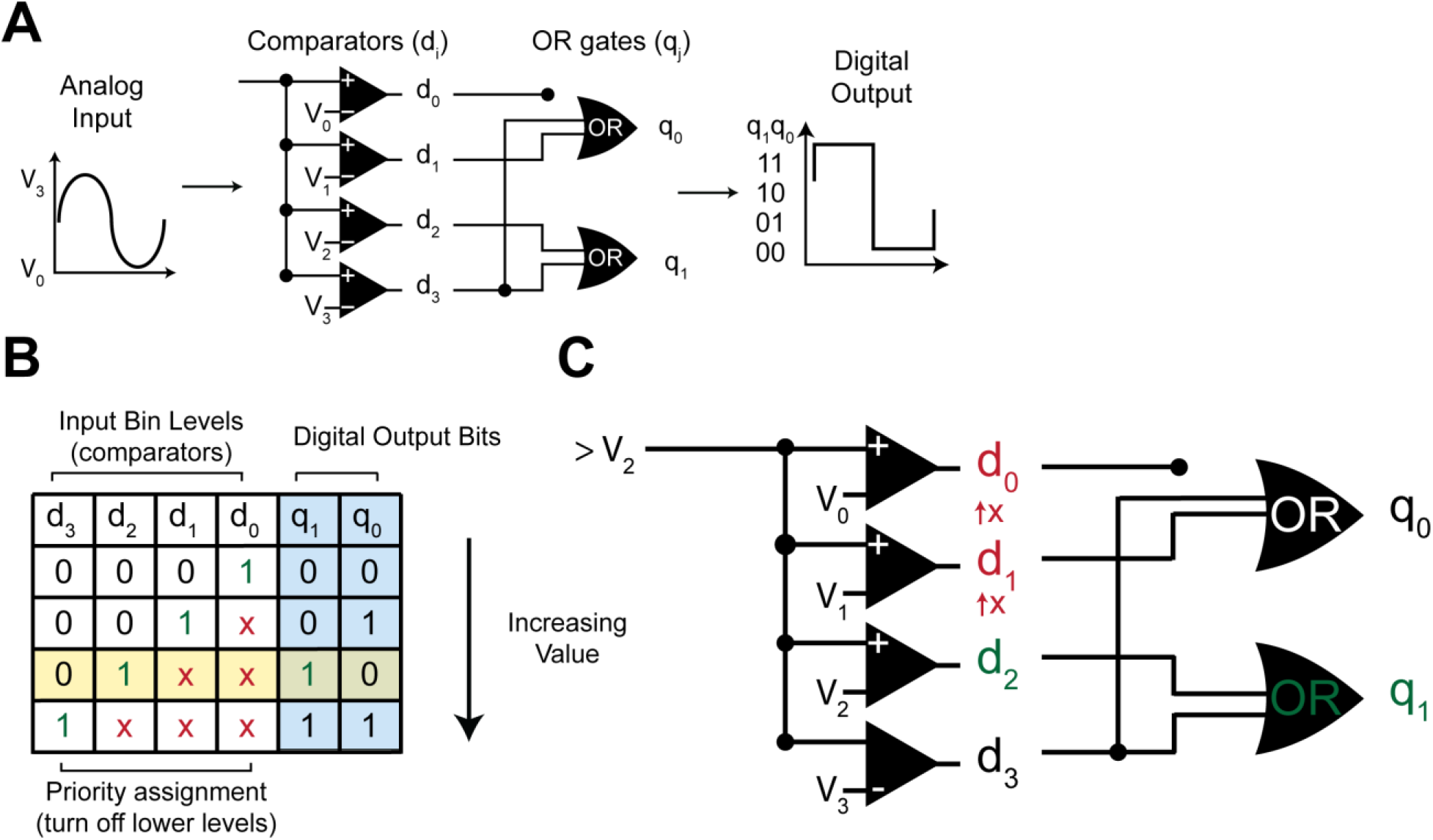
Inputs and outputs of a 4-2 bit ADC. **(A)** Circuit diagram of a flash ADC. **(B)** Truth table. Inputs activate continuous subsets of the biocomparators (d_0_ to d_3_). An input which activates a biocomparator produces a value of 1. To give this input priority, all biocomparators below are turned off, signified by a value of x. Digital output bits are colored blue and correspond to the 2-bit output of the ADC. **(C)** Logic circuit diagram for one example input/output case through a 4-2 bit ADC. Input signal > V_2_ turns on d_0_—d_2_, but priority is given to d_2_, which only turns on bit q_1_, producing the output 10.

**Fig. S3.**
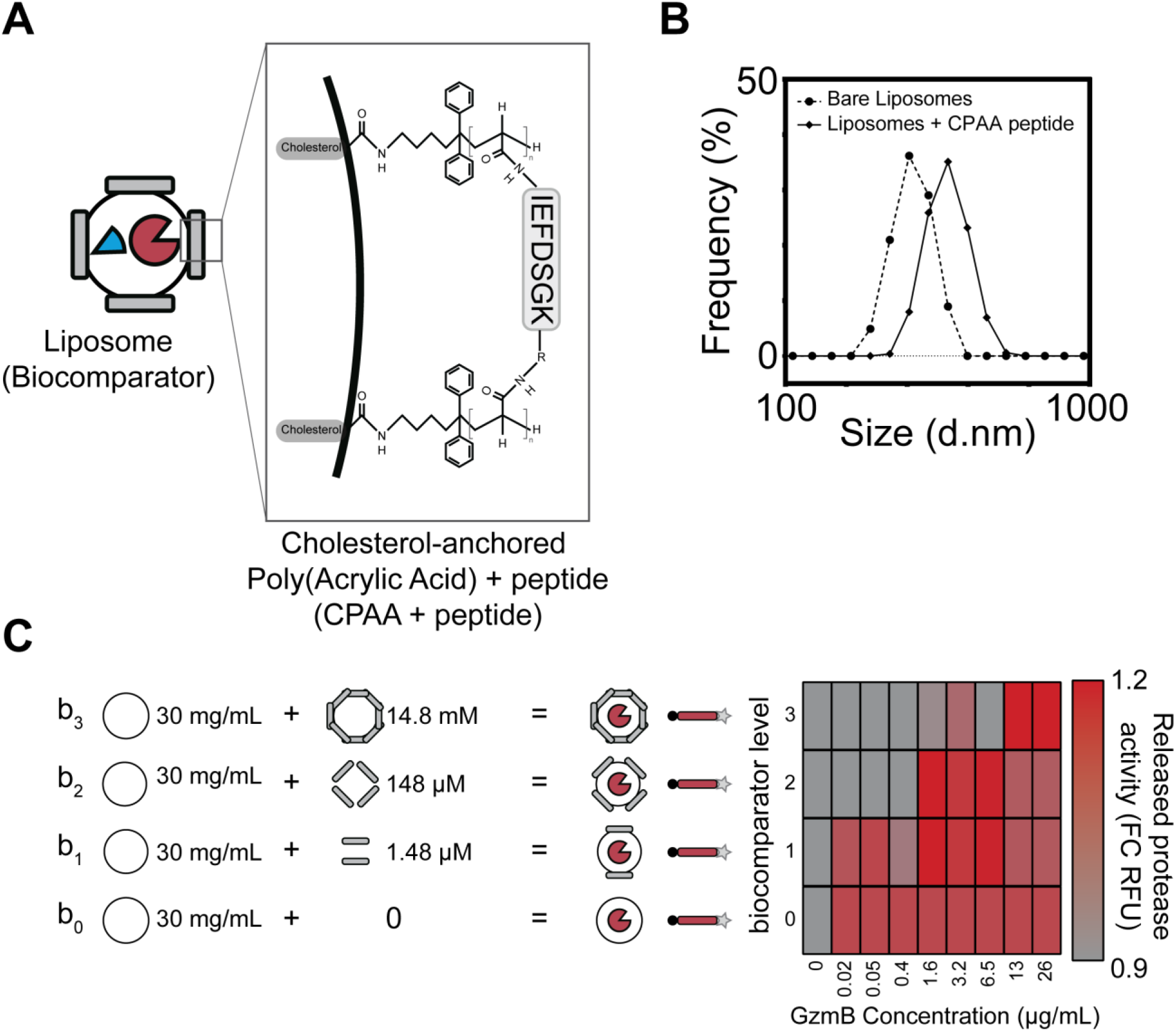
Peptide caged liposome synthesis and characterization. **(A)** Graphic of cholesterol-anchored poly(acrylic acid) (CPAA) embedded in liposome membrane, crosslinked by amine terminated peptides. Carboxylic acid side groups on poly(acrylic acid) are activated by EDC*MeI. N-terminal amine and C-terminal amine (from lysine side chain) act as primary amines to react with activated CPAA side chains. **(B)** DLS size measurement of liposomes before and after peptide cage construction. Increase in average hydrodynamic radius from 264.0 nm (bare liposomes) to 344.3 nm (Liposomes + CPAA peptide). **(C)** Heat map showing concentration of GzmB required to unlock each level. Signal is measured via released protease cutting substrate, normalized to the negative control (0 ug/mL signal protease).

**Fig. S4.**
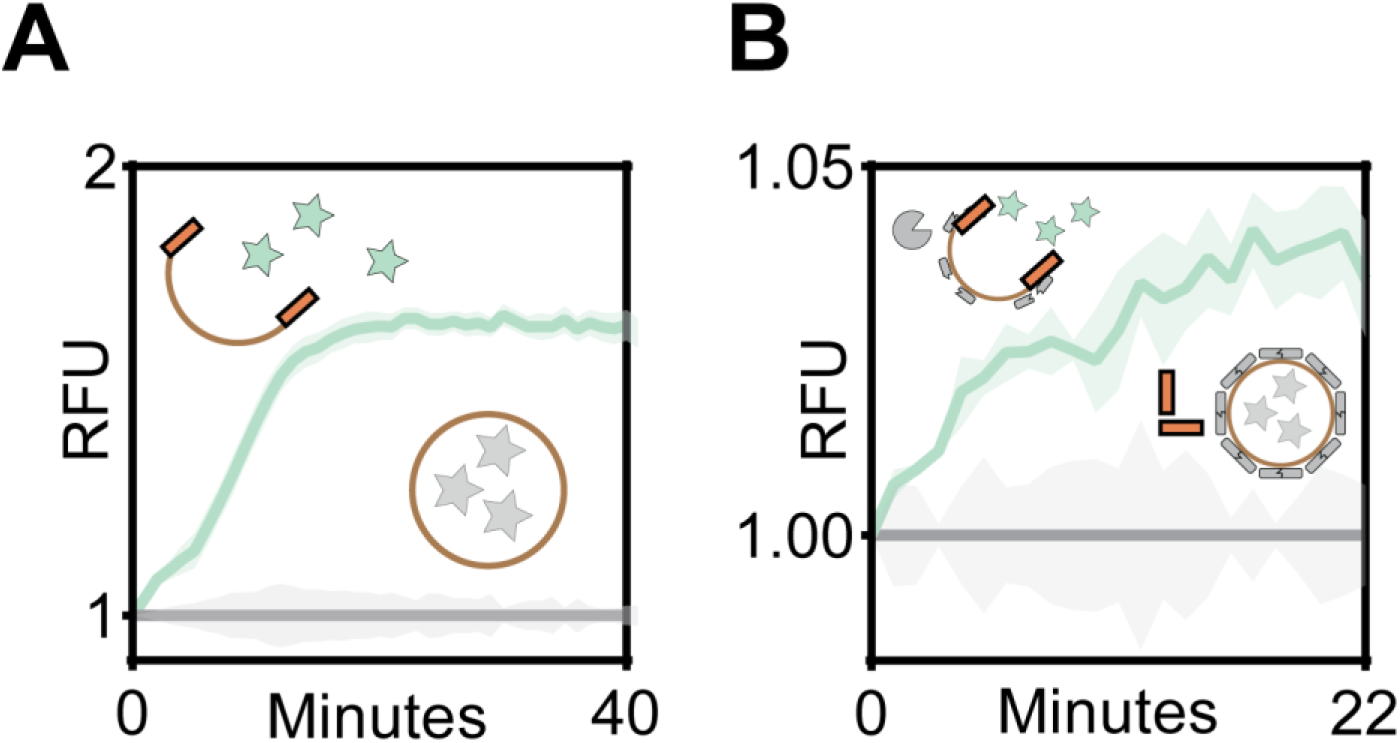
Lipase acts as a Buffer gate. **(A)** Phospholipase C-triggered release of FITC contained in liposomes. Negative control contains liposome only and no lipase. **(B)** Phospholipase-C and signal protease GzmB triggered release of FITC. Negative control contains lipase, but no signal protease. Line shading represents standard deviation (n = 3).

**Fig. S5.**
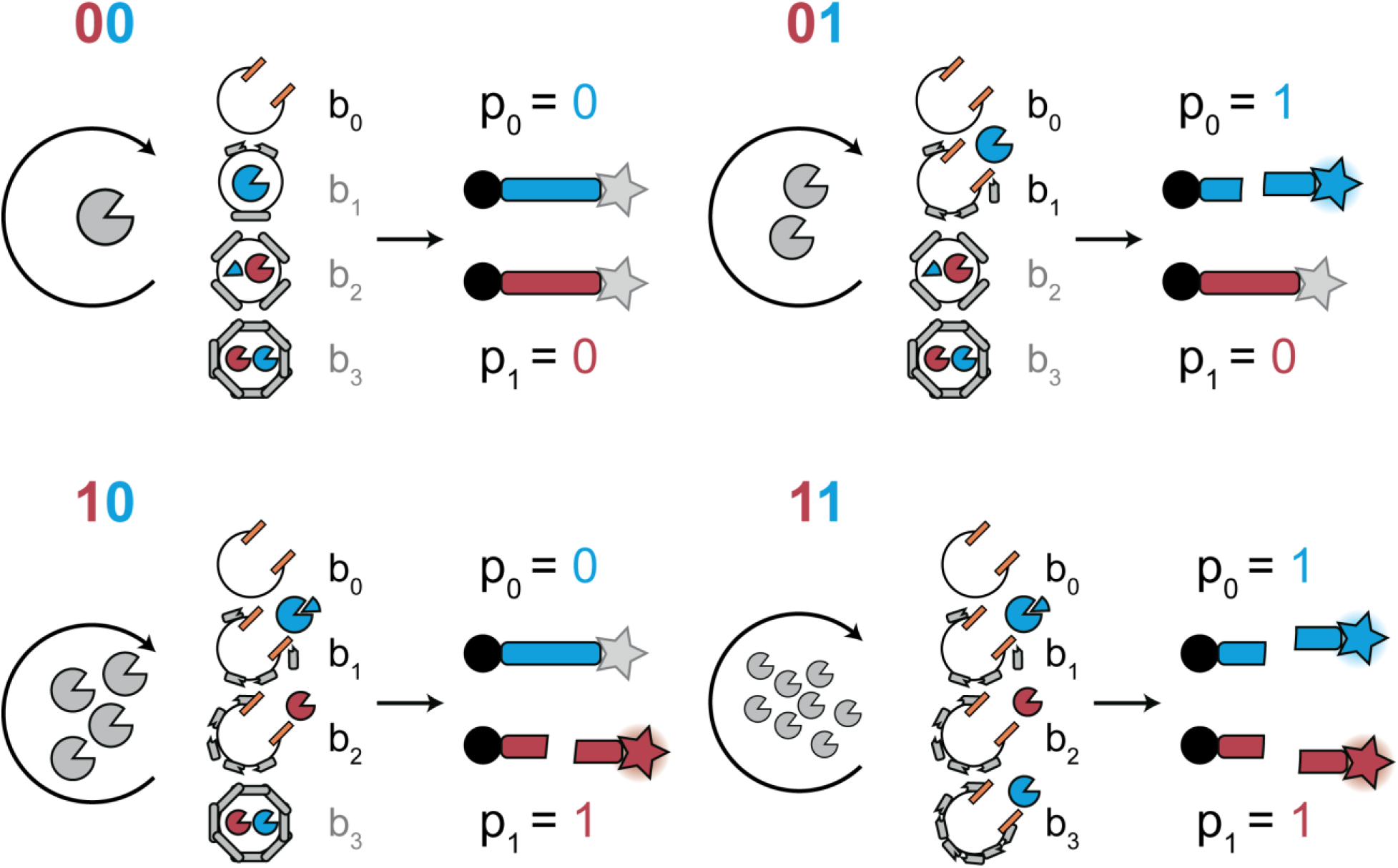
All possible digital outputs of the bioADC. Schematic of all 4 possible signal conversions in the biological four-two bit analog-to-digital converter. The signal protease (grey, GzmB), cleaves the peptide cage surrounding biocomparators. Higher activity levels of GzmB result in more biocomparator levels being unlocked (b_0_ to b_3_). Lipase (orange rectangle) is co-incubated with the bioADC such that all exposed biocomparators are fully opened via degradation of liposome by phospholipase C. Released signal converter proteases and priority encoding inhibitors interact to produce one digital signal. This signal interacts with OR gates to produce “high”, or 1, values for the correct binary digits.

**Fig. S6.**
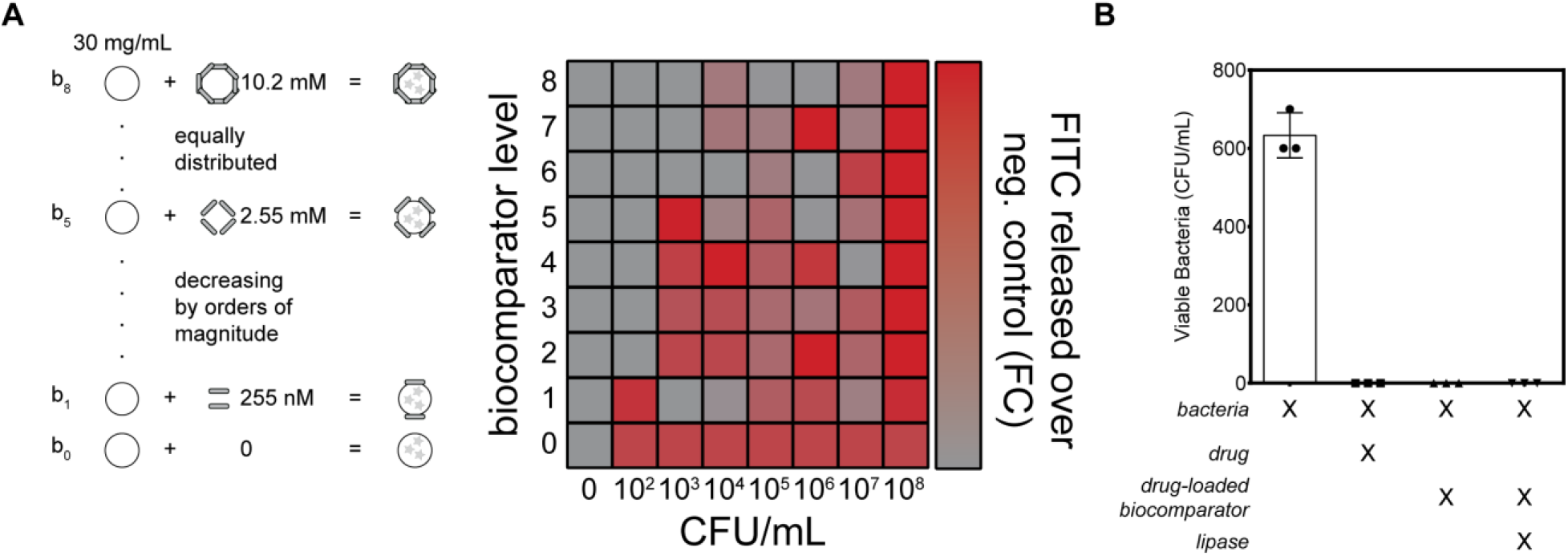
Biocomparator and OR gate interface with bacteria. **(A)** Unlocking peptide caged liposomes with increasing peptide crosslinking densities (levels 0 – 8). Eight levels of increasing peptide cage crosslinking were used to determine the number of bacteria required to unlock each level. Increased concentration of input protease (OmpT, from *E. coli*) leads to more unlocked levels. **(B)** Bacterial cytotoxicity measurement of drug-loaded liposomes. Conditions are moving from left to right: Bacteria only control, bacteria plus free drug, bacteria plus drug-loaded liposome without lipase, and bacteria plus drug-loaded liposome with lipase. Samples were incubated with bacteria at 37C for eight hours and plated. CFU were quantified to estimate bacteria viability. Error bars are plotted as standard deviation (n = 3).

**Fig. S7.**
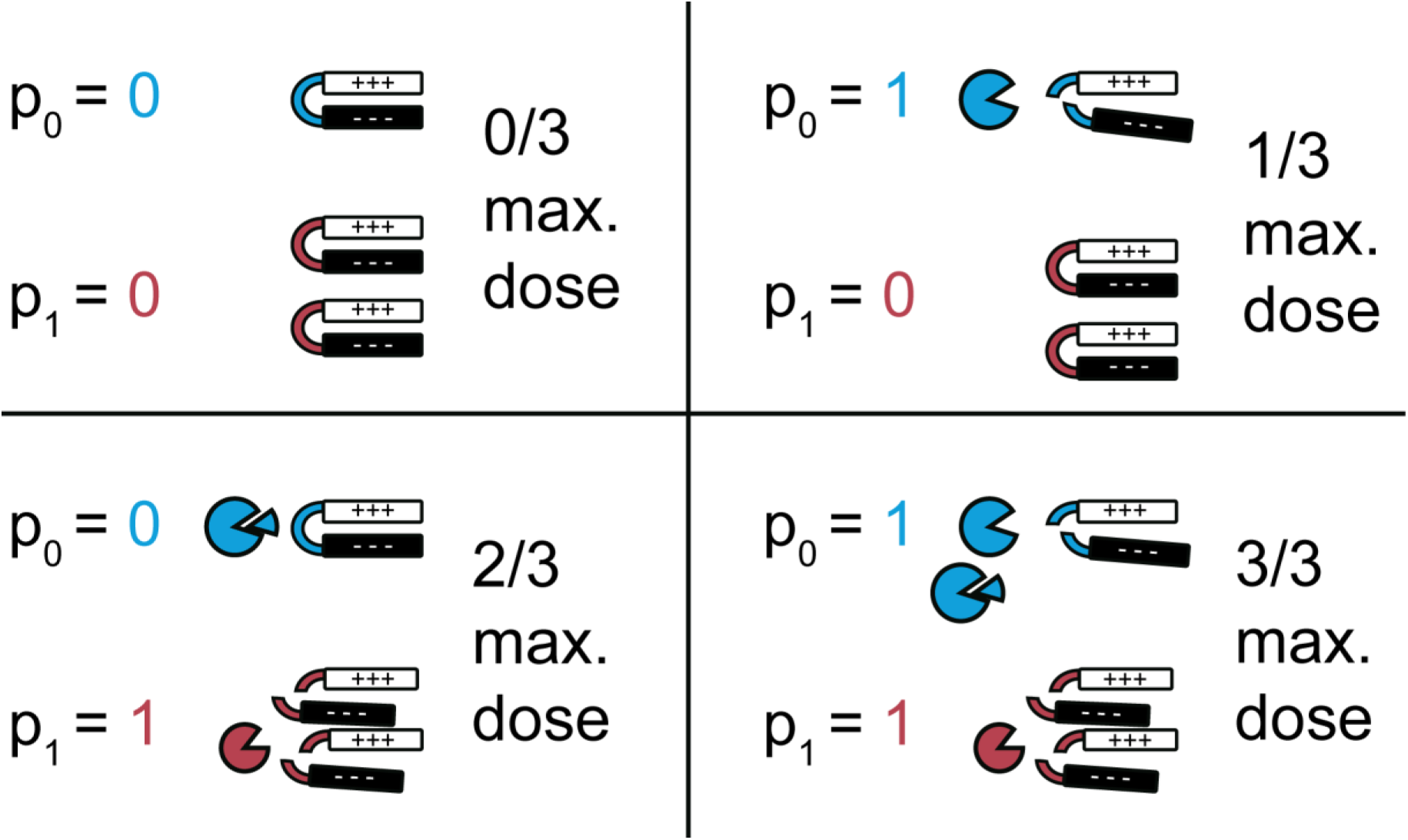
All possible drug doses from bioprogram. The OR gate 0 linked to bit p_0_ outputs 1/3 of the available AMP dose, whereas the OR gate 1 linked to bit p_1_ outputs 2/3 of the available AMP dose. This translates each digital output to a drug dose increasing by units of 1/3 the total dose.

**Fig. S8.**
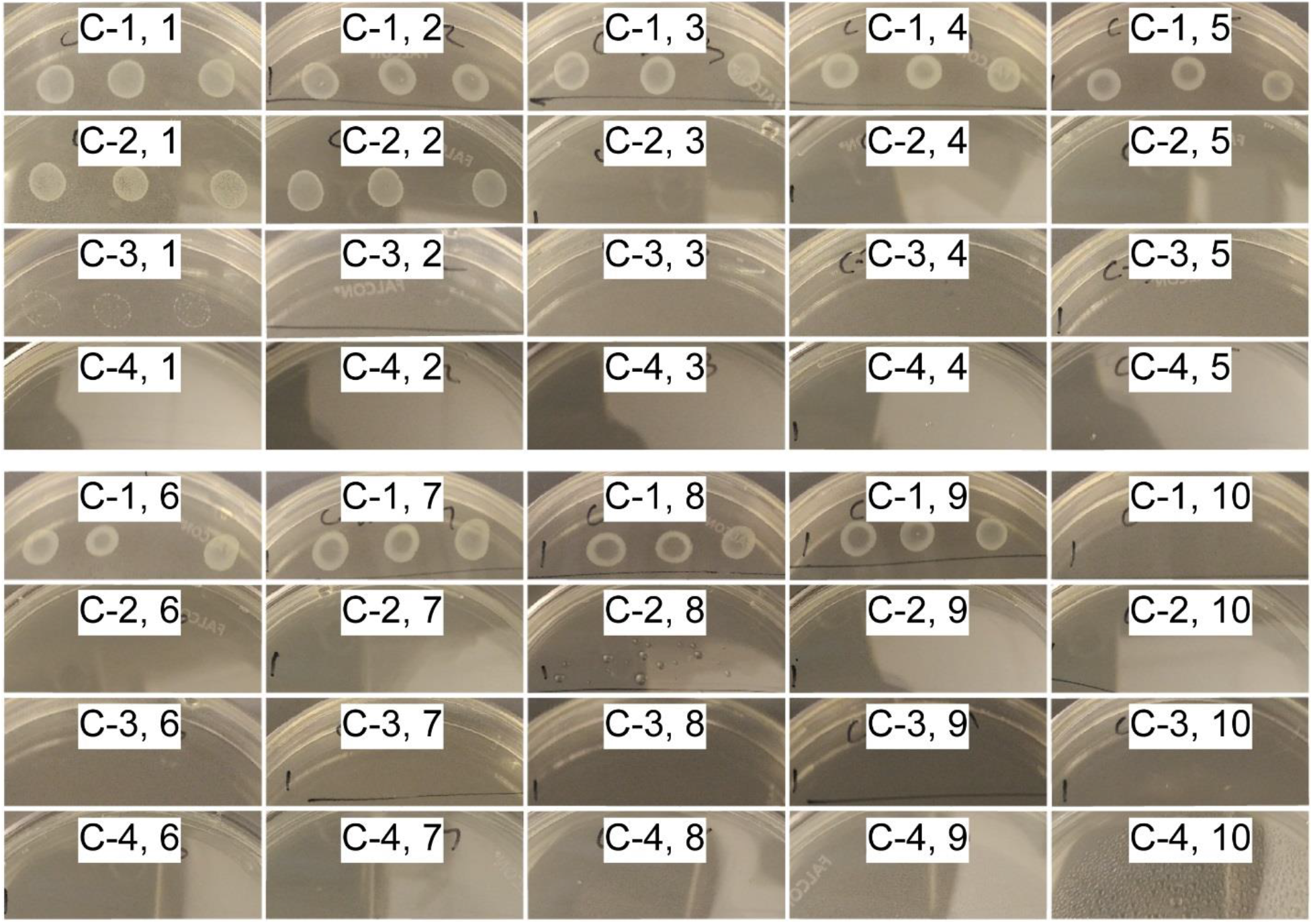
Photos of bacterial plates post-incubation with bioprograms. Photos of bacterial plates used for quantification of cytotoxicity in Fig 3E. C-1 indicates the single, empty comparator bioprogram control, which is defined as 100% cell viability, or 0% cytotoxicity. No colony growth is defined as 0% cell viability, or 100% cytotoxicity. Plates are labeled C-*n X*, where *n* corresponds to the number of biocomparators p resent in the program combined with infected blood and *X* corresponds to the concentration of bacteria present. (10 = 10^0^ CFU/mL, 9 = 10^1^ CFU/mL, 8 = 10^2^ CFU/mL, etc.).

**Fig. S9.**
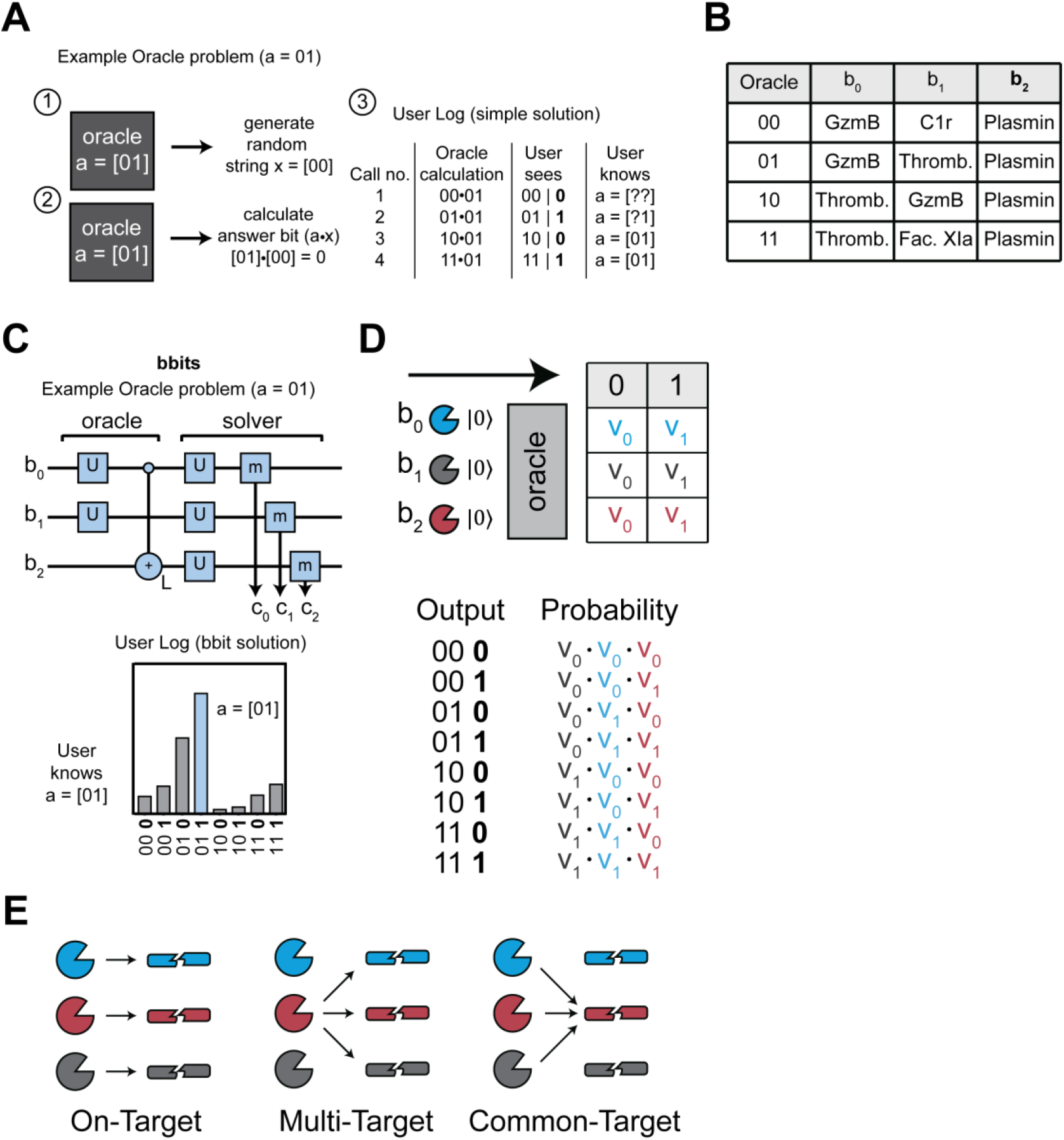
Implementing the oracle problem with protease bbits. **(A)** For example, Learning Parity with Noise (LPN) is a general inference problem where an oracle (i.e., a black box) is hiding a string of bits, which should be learned with the smallest number of oracle queries. Upon each query, the oracle (1) generates a random string of bits and (2) takes the dot product of the hidden and random strings, which produces the answer bit. Due to the definition of the dot product, each time the answer bit takes on a value of 1, the oracle reveals which digits in the hidden string may also take the value 1 (i.e., which digits in the hidden string exhibit parity with the answer bit), eventually yielding the one correct string. However, in the presence of noise, or uncertainty, the oracle makes mistakes that result in misinformation, making it more difficult to infer the value of the hidden string. A quantum algorithm accounts for this uncertainty by combining the information from all oracle queries to calculate the probabilities that each possible string is the correct one and choosing the string with the highest probability^42^. **(B)** Table of protease bits used to implement all 4 possible biological oracles. **(C)** Workflow involved in solving the same example of the two-bit oracle problem using quantum-inspired bbits. Top depicts the biological analog of a quantum score with the orientation of gates and bbits required to simulate and then solve the Oracle problem. **(D)** Calculating probabilities of all 8 possible outputs from the 2-bit oracle problem are calculated from multiplying relative cleavage velocities (v_n_) of each bbit. **(E)** Schematic demonstrating on-target, multi-target, and common-target protease activity.

**Fig. S10.**
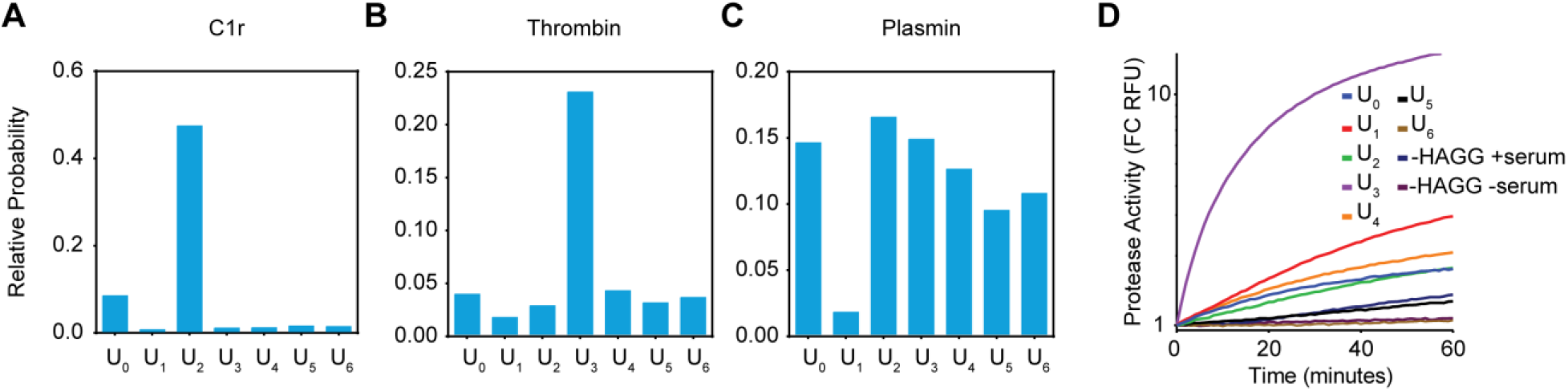
Characterizing protease specificity towards U gates. **(A-C)** Cleavage assay measuring complement (C1r) and coagulation (Thrombin, Plasmin) protease specificity towards U-gates U_0_ to U_6_, panels A to G, respectively. **(D)** Human serum complement activation assay to measure specificity of proteases in the classical complement cascade towards U-gates U_0_ to U_6_.

**Fig. S11.**
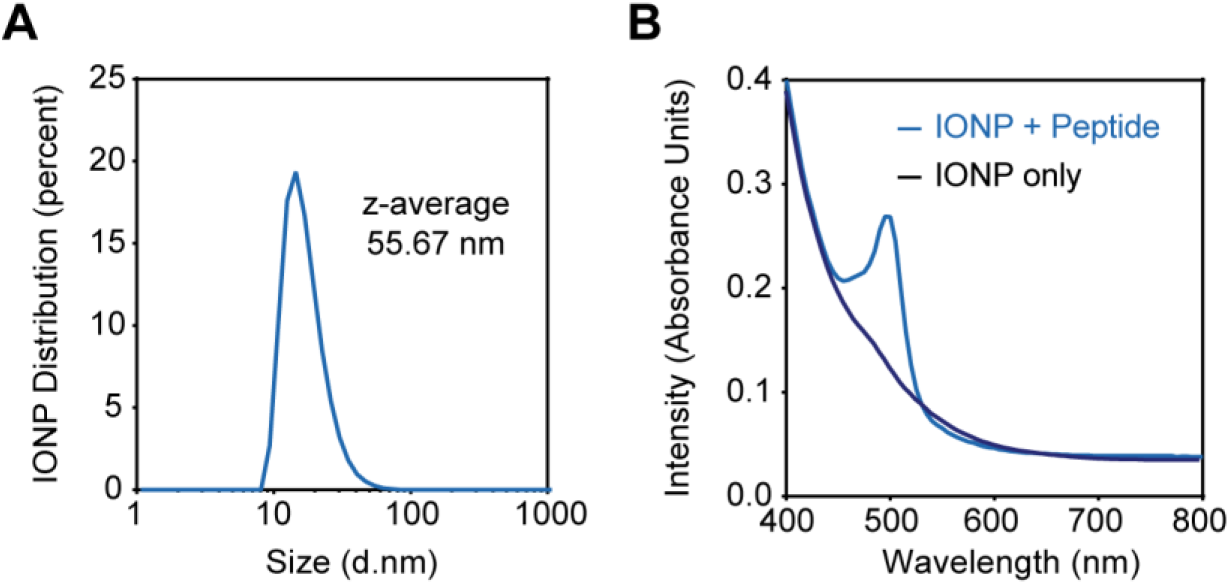
Characterizing Iron Oxide Nanoparticles for *in vivo* administration. **(A)** Dynamic Light Scattering (DLS) measurement of nanoparticle size distribution. **(B)** Absorbance spectra for iron oxide nanoparticles (IONP) only and IONP conjugated to a substrate, measured in 5 nm steps from 400 to 800 nm.

**Fig. S12.**
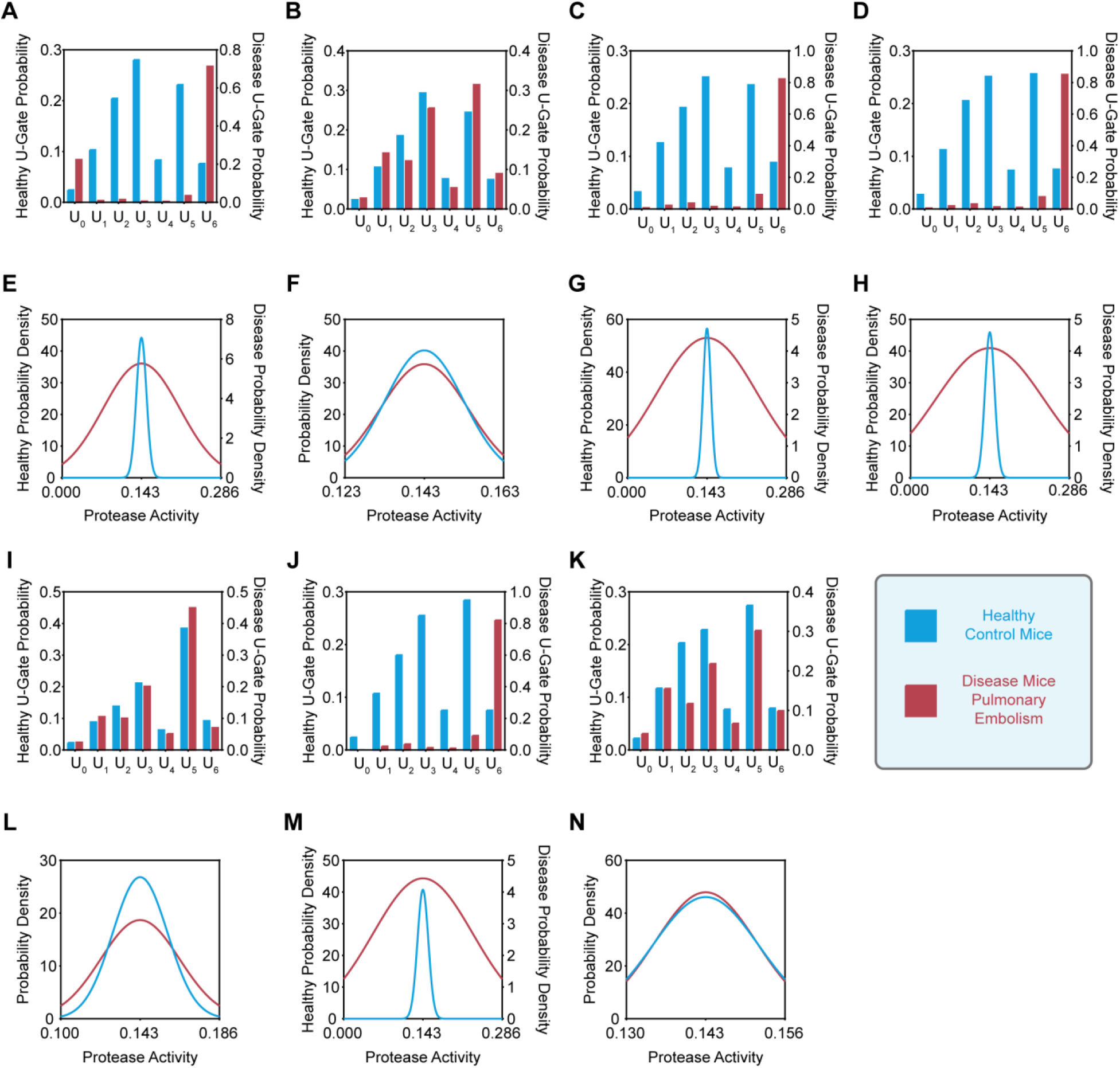
Extracting protease activity probabilities with biological “oracle” algorithm. **(A— D, I—K)** Relative U-gate probabilities measured via urinalysis of the seven mice before (blue) and after (red) onset of pulmonary embolism (Injection 1). **(E—H, L—N)** Extraction of protease activity probability distribution profiles before (blue) and after (red) onset of pulmonary embolism.

**Table S1.**
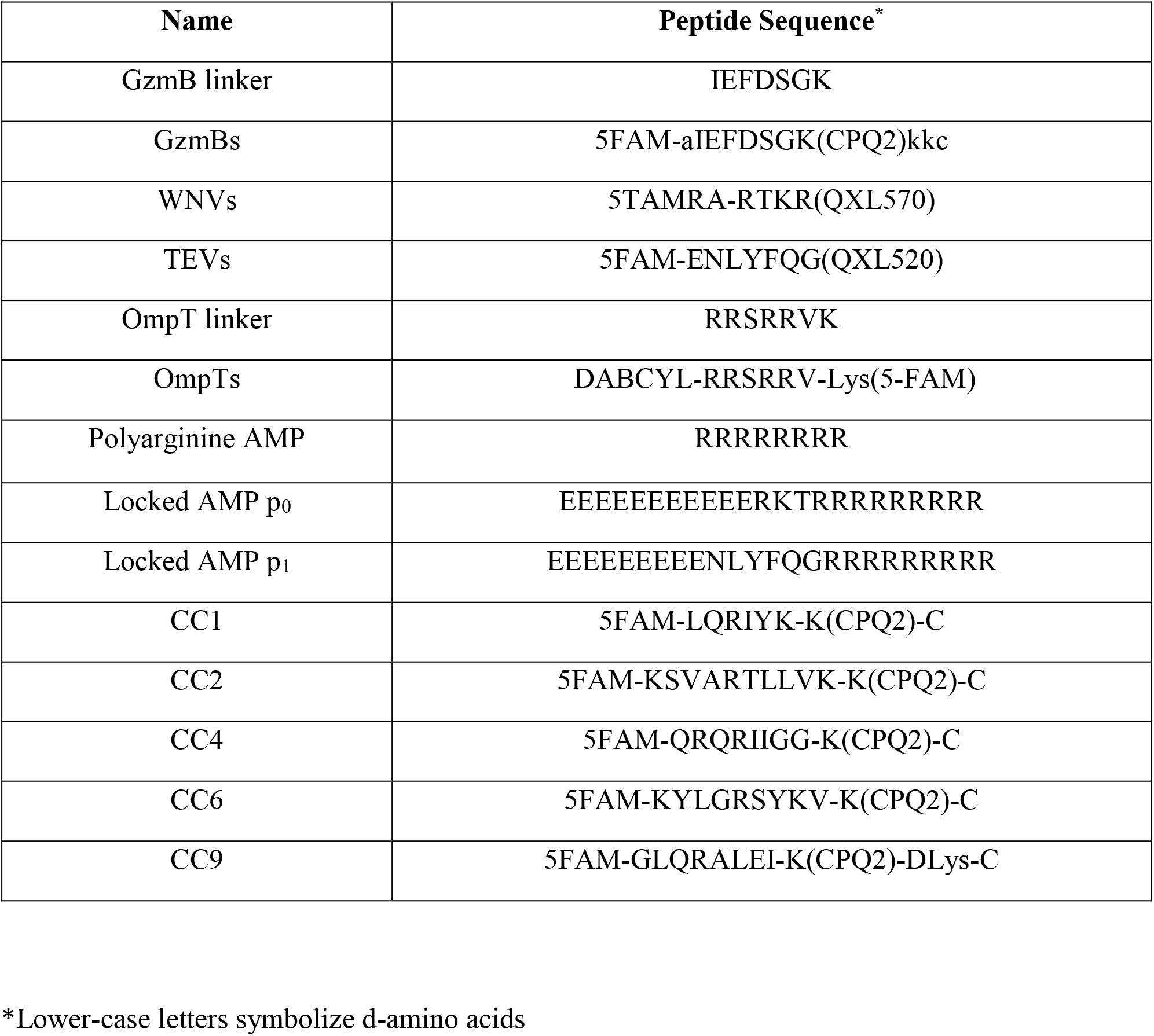

| **Name** | **Peptide Sequence\*** |
| --- | --- |
| GzmB linker | IEFDSGK |
| GzmBs | 5FAM-aIEFDSGK(CPQ2)kkc |
| WNVs | 5TAMRA-RTKR(QXL570) |
| TEVs | 5FAM-ENLYFQG(QXL520) |
| OmpT linker | RRSRRVK |
| OmpTs | DABCYL-RRSRRV-Lys(5-FAM) |
| Polyarginine AMP | RRRRRRRR |
| Locked AMP p<sub>0</sub> | EEEEEEEEEEERKTRRRRRRRRR |
| Locked AMP p<sub>1</sub> | EEEEEEEEENLYFQGRRRRRRRRR |
| CC1 | 5FAM-LQRIYK-K(CPQ2)-C |
| CC2 | 5FAM-KSVARTLLVK-K(CPQ2)-C |
| CC4 | 5FAM-QRQRIIGG-K(CPQ2)-C |
| CC6 | 5FAM-KYLGRSYKV-K(CPQ2)-C |
| CC9 | 5FAM-GLQRALEI-K(CPQ2)-DLys-C |

\*Lower-case letters symbolize d-amino acids

**Table S2.**

| **Protease Name** | **Abbreviation** | **Substrate(s)** |
| --- | --- | --- |
| Granzyme B | GzmB | IEFDSG, IEFDSGK |
| West Nile Virus Protease | WNVp | RTKR, inhibitor: undeca-D-ArgNH<sub>2</sub> |
| Tobacco Etch Virus Protease | TEVp | ENLYFQG |
| Outer Membrane Protein T | OmpT | RRSRRV |
| Complement protease C1r | C1r | QRQRIIGG, LQRIYK |
| Thrombin | Throm. | KSVARTLLVK, KYLGRSYKV |
| Plasmin | Plasm. | GLQRALEI, KYLGRSYKV |
| Factor XIa | Fac. XIa | LQRIYK, KYLGRSYKV |

**Table S3.**

| **Substrate Name** | **Peptide sequence (N terminus on left)** | **Modifications** |
| --- | --- | --- |
| U<sub>0</sub> | e(*aa)(*aa)ndneeGFFsAr(ANP)K(5-FAM)GGLQRIYKC | 1st *aa= Gly(13C2); 2nd *aa=Val(U13C5,15N) |
| U<sub>1</sub> | eG(*aa)ndneeGF(*aa)s(*aa)r(ANP)K(5-FAM)GGKSVARTLLVKC | 1st *aa= Val(U13C5,15N); 2nd *aa=Phe(15N); 3rd *aa=Ala(15N) |
| U<sub>2</sub> | e(*aa)(*aa)ndneeGFFs(*aa)r(ANP)K(5-FAM)GGQRQRIIGGC | 1st *aa= Gly(U13C2,15N); 2nd *aa=Val(15N); 3rd *aa=Ala (U13C3,15N) |
| U<sub>3</sub> | e(*aa)Vndnee(*aa)FFs(*aa)r(ANP)K(5-FAM)GGKYLGRSYKVC | 1st *aa= Gly(13C2); 2nd *aa=Gly(13C2); 3rd *aa=Ala(U13C3,15N) |
| U<sub>4</sub> | eGVndnee(*aa)(*aa)Fs(*aa)r(ANP)K(5-FAM)GGGLQRALEIC | 1st *aa=Gly(U13C2,15N); 2nd *aa=Phe(15N); 3rd *aa=Ala(U13C3,15N) |
| U<sub>5</sub> | e(*aa)(*aa)ndnee(*aa)(*aa)(*aa)s(*aa)r(ANP)K(5-FAM)GGKTTGGRIYGGC | 1st *aa=Gly(13C2); 2nd *aa=Val(U13C5,15N); 3rd *aa=Gly(U13C2,15N); 4th *aa=Phe(15N); 5th *aa=Phe(15N); 6th *aa=Ala(15N); still include ANP and K5-FAM |
| U<sub>6</sub> | eG(*aa)ndnee(*aa)(*aa)Fs(*aa)r(ANP)K(5-FAM)GGQARGGSC | 1st *aa=Val(U13C5,15N); 2<sup>nd</sup> *aa=Gly(U13C2,15N); 3rd *aa=Phe(15N); 4th *aa=Ala(U13C3,15N) |

